# Structure and mechanism of the HSV-1 origin-binding protein UL9

**DOI:** 10.1101/2025.10.31.685793

**Authors:** Cuiqing Huang, Haiqiang Wu, Jinmiao Song, Xinzheng Zhang, Jun Ma

**Author notes:** These authors contributed equally: Cuiqing Huang, Haiqiang Wu. Correspondence: Jun Ma, Xinzheng Zhang.

## Abstract

The herpesvirus DNA replication machinery comprises a battery of viral enzymes that orchestrate viral genome synthesis. In herpes simplex virus type 1 (HSV-1), the machinery consists of seven essential components, including the origin-binding protein UL9, the single-stranded DNA (ssDNA)-binding protein ICP8, the heterodimeric DNA polymerase complex UL30–UL42, and the heterotrimeric helicase–primase complex UL5–UL8–UL52. UL9, a superfamily 2 (SF2) helicase, functions as a dimer that specifically recognizes replication origins and unwinds duplex DNA to initiate replication. Furthermore, UL9 recruits the replication machinery through interactions with viral components and engages cellular proteins that regulate its function. However, the molecular mechanisms underlying UL9’s multifunctionality remain incompletely understood due to the lack of structural information. Here, we present cryo-electron microscopy structures of UL9 in both apo and DNA-bound states. Together with biochemical and enzymatic assays, we elucidate the molecular basis of UL9 dimerization, origin recognition and allosteric regulation by ICP8.

## Introduction

The *Herpesviridae* family comprises a large group of enveloped viruses with relatively large, complex double-stranded DNA genomes. To date, over 100 herpesviruses with a wide range of vertebrate hosts have been recognized and classified into three subfamilies (α-, β- and γ-herpesviruses) based on their biological properties and tissue tropism. Notably, nine herpesviruses are pathogenic in humans and cause numerous human diseases; these include α-herpesviruses (herpes simplex virus types 1 and 2 (HSV-1, HSV-2), varicella-zoster virus (VZV)); β-herpesviruses (human cytomegalovirus (HCMV), human herpesvirus 6A/B (HHV-6A, HHV-6B), human herpesvirus 7 (HHV-7)); γ-herpesviruses (Epstein-Barr virus (EBV) and Kaposi’s sarcoma-associated herpesvirus (KSHV/HHV-8))[1]. Over 90% of the global population is seropositive for one or more of these viruses. Herpesviruses establish lifelong latency in infected persons and periodically reactivate during the episodes of immunosuppression or physiological stress, with no therapeutic approaches currently available to achieve complete viral eradication[2]. To date, there are limited medical interventions for specific herpesviruses, including two approved vaccines Shingrix and Zostavax for VZV[3], and antiviral drugs such as Acyclovir, Valacyclovir, Famciclovir and Pritelivir effective against HSV-1 and VZV[4]. For other herpesviruses, particularly oncogenic pathogens EBV and KSHV, effective antiviral drugs or vaccines remain unavailable. Furthermore, current antivirals carry risks of side effects and drug resistance[4-6]. Consequently, the development of novel medical treatments against herpesviruses is essential, with viral DNA replication representing a particularly promising target[7, 8].

Herpesvirus DNA replication occurs in the nucleus of the host cell, carried out by a sophisticated multienzyme DNA replication machinery[9-11]. In HSV-1, this machinery comprises seven core components, including the replication origin-binding protein (OBP) UL9, single-stranded DNA (ssDNA)-binding protein ICP8, the heterodimeric DNA polymerase complex UL30/UL42, and the heterotrimeric helicase-primase complex UL5/UL8/UL52. The DNA polymerase complex possesses two functionally distinct subunits: the catalytic subunit UL30 is a B-family DNA polymerase harboring both 5’-3’ polymerase activity and 3’-5’ proofreading exonuclease activity, and the processivity factor UL42 helps polymerase maintain high processivity. Within the helicase-primase complex, the helicase subunit UL5 is responsible for double-stranded DNA (dsDNA) unwinding during elongation, the primase subunit UL52 synthesizes short RNA primers for DNA polymerase, and the primase-helicase associated factor UL8 coordinates the activities of UL5 and UL52. The ssDNA-binding protein ICP8 is a multifunctional protein that prefers to bind ssDNA in a non-sequence-specific and cooperative manner while preventing it from nuclease degradation and annealing. Additionally, ICP8 stimulates the helicase activity of UL9, modulates DNA polymerase processivity, and regulates helicase-primase complex function through direct interactions with UL9, UL42 and UL8[9-11], respectively. The origin-binding protein UL9 mainly orchestrates replication initiation through sequence-specific recognition of origin sites and subsequently unwinds the genomic DNA with the help of ICP8 to initiate DNA replication[9-20]. Furthermore, UL9 functions as a molecular hub, recruiting the replication machinery via interactions with ICP8[17, 19, 21, 22], UL42[23-25] and UL8[26], while engaging cellular proteins[11] such as topoisomerase I[22], hTid-1[27], and heat shock proteins[28] to regulate its own functions. These multifaceted roles establish UL9 as the central coordinator of herpesvirus DNA synthesis.

As a member of the SF2 helicase superfamily, HSV-1 UL9 has been reported to function as a homodimer under physiological conditions[15, 29], with each protomer comprising an N-terminal helicase domain and a C-terminal domain (Fig. 1a). Despite its essential role, the molecular mechanism of UL9 remains elusive due to the lack of structural information. In this study, we present the structures of HSV-1 UL9 in both apo and DNA-bound states, determined by cryo-electron microscopy (cryo-EM) single-particle analysis. Combined with *in vitro* biochemical and enzymatic assays, we elucidate the molecular basis of UL9’s dimerization, origin recognition and activity regulation by ICP8 interaction. These findings not only advance our understanding of the mechanism of HSV-1 replication initiation but also provide potential new targets for the development of anti-herpesviral drugs.

**Figure 1.**
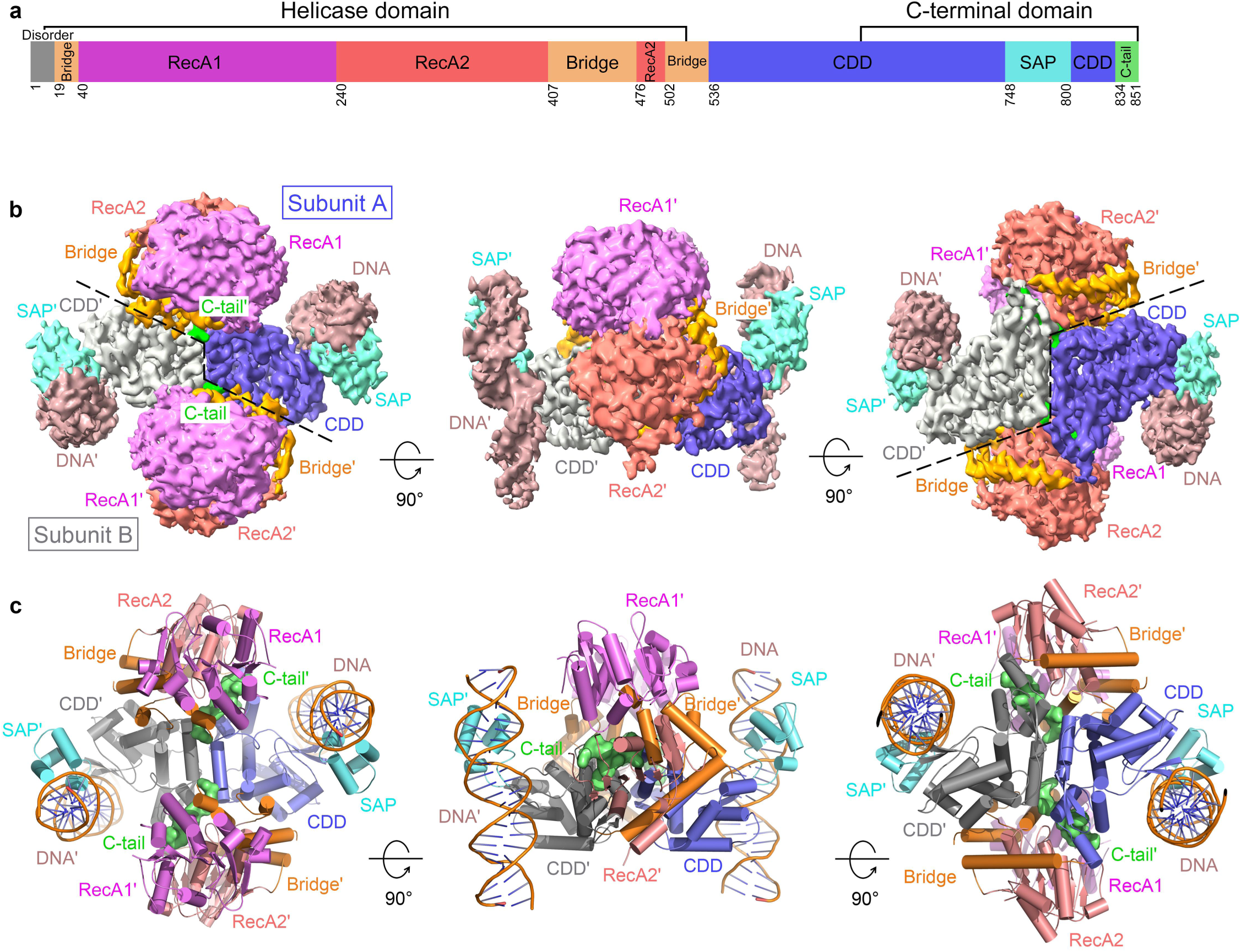
Cryo-EM structure of the UL9-DNA complex. **a** Domain architecture of HSV-1 UL9 protein, with the N-terminal helicase domain and C-terminal domain indicated. The RecA1, RecA2, Bridge, CDD, SAP and C-tail subdomains or regions are shown in violet, salmon, orange, slate, aquamarine and lime, respectively. Inter-subdomain boundaries are marked with residue numbers. **b** Three orthogonal views of the cryo-EM density map for the UL9-DNA complex, colored according to the domain coloring scheme in (**a**). **c** Corresponding orthogonal views of the atomic model presented in cartoon representation, colored identically to panel **b**.

## Result

### Cryo-EM structure determination of HSV-1 UL9

To determine the structure of HSV-1 UL9, the full-length protein (UL9^FL^) was expressed in insect cells and purified to homogeneity using a tandem two-step purification strategy (Supplementary Fig. 1). The UL9-DNA complex was prepared by incubating affinity-purified UL9 with 56-bp dsDNA containing *OriS* Box I and II at a molar ratio of 1:1.5, followed by gel filtration (Supplementary Fig. 1). Using cryo-EM single-particle analysis, the structures of apo-UL9 and the UL9-DNA complex were resolved at global resolutions of 4.1 Å and 3.7 Å, respectively (Fig. 1, Supplementary Figs. 2 and 3, and Supplementary Table 1). Due to the higher resolution of the map, the model of the UL9-DNA complex was first built with the aid of Alphafold2 prediction[30]; this model was primarily used for subsequent structural analyses. The final model of the UL9-DNA complex comprises residues 19-843 and two DNA duplexes, although flexible loops (residues 135-146, 267-283, and 431-456) remain unresolved (Fig. 1c and Supplementary Fig. 4a). The DNA-binding region (residues 748-799) and the bound DNA exhibited significant flexibility, limiting the local resolution to approximately 4.5 Å (Supplementary Fig. 2g). Consequently, we modeled this domain and a 23-bp DNA segment by rigid-body fitting of the AlphaFold-predicted structure into the density map. While the specific nucleotide sequence registration in the model remained undetermined, preventing analysis of sequence-specific recognition mechanisms, the overall architecture is well-defined. In contrast to the DNA-bound state, residues 748-799 in the apo-UL9 structure were disordered and could not be modeled due to a lack of defined density (Supplementary Fig. 4b).

### Architecture of the UL9 dimeric complex

Consistent with previous studies demonstrating that UL9 functions as a dimer in solution and *in vivo*[15, 29], both our apo and DNA-bound structures reveal a C2-symmetric dimer. The UL9-DNA complex measures approximately 115 × 110 × 65 Å (Supplementary Fig. 4a). In the apo state, the DNA-binding elements are disordered, resulting in a more slender yet compact structure core (Supplementary Fig. 4b). Despite the compact dimeric core, individual protomers adopt an extended architecture (Figs. 2a, b) that clearly delineates the N-terminal helicase domain (residues 19-535) and C-terminal domain (CTD, residues 536-851). Dimerization is mediated primarily by closely apposed CTDs, which form a rigid central core flanked by two helicase domains; notably, the helicase domains do not interact with one another. Apart from the ordering of the DNA-binding elements, DNA binding induces no major global conformational changes (Supplementary Fig. 4c).

**Figure 2.**
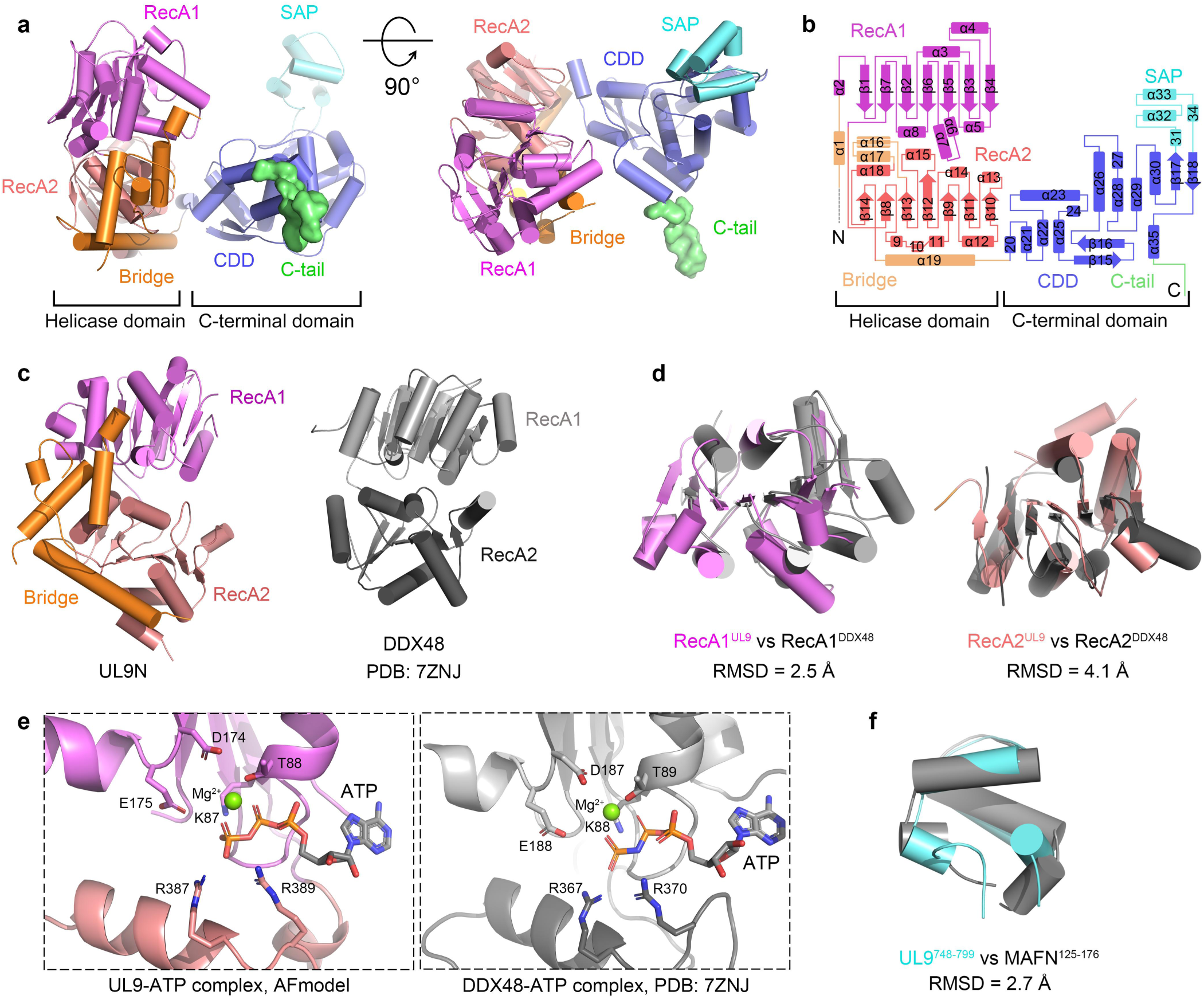
Structural analysis of UL9 protomer. **a** Two orthogonal views of the UL9 protomer structure. **b** Two-dimensional topology diagram of UL9. **c** Structural comparison of UL9 helicase domain with its structural homolog DDX48 identified by the Dali server. **d** Structural superposition of UL9 RecA1 and RecA2 subdomains with corresponding subdomains in DDX48. **e** Key residues involved in ATP binding and hydrolysis in UL9 are identical to those in its structural homolog DDX48. The UL9-ATP complex model was predicted with AlphaFold. **f** Structural superposition of UL9 SAP domain with its homolog MAFN identified by the Dali server. All structural representations throughout the panels follow the color scheme in Figure 1a, with domains labeled and homologous proteins (DDX48 and MAFN) shown in gray. RMSD values for structural superpositions in panels **d** and **f** are provided.

### Structural details of UL9 helicase domain

Structural homology analysis of the N-terminal helicase domain using the Dali server[31] identified DDX48[32] (PDB: 7ZNJ, Z=18.9, RMSD=3.8 Å) (Fig. 2c), DDX17[33] (PDB: 6UV3, Z=18.7, RMSD=3.9 Å), and Vasa[34] (PDB: 4D26, Z=15.9, RMSD=3.9 Å) as top hits, all of which belong to the DExD/H-box SF2 helicase superfamily. Consistent with other SF2 members, UL9’s helicase domain can be subdivided into two RecA-like subdomains: RecA1 (residues 40-242) and RecA2 (residues 243-406 and 476-501) (Figs. 1a and 2a-d). Superposition of these subdomains with their counterparts in DDX48 (RecA1, residues 23-242; RecA2, residues 243-404) revealed root-mean-square deviation (RMSD) values of 2.5 Å and 4.1 Å (Fig. 2d), respectively, confirming that UL9 retains the conserved RecA-like architecture characterized by seven parallel β-strands. Furthermore, structural alignment of an AlphaFold-predicted UL9–ATP model with DDX48 indicates that key residues involved in ATP binding and hydrolysis within RecA1 are identical (Fig. 2e). However, the UL9 helicase domain also contains additional structural elements (residues 19-39, 407-475 and 502-535) outside the conserved RecA1 and RecA2 subdomains (Fig. 2d). The discontinuous segments assemble into a distinct structural feature spanning RecA1 and RecA2, which we term the **Bridge motif** (Fig. 2a-c). As detailed below, the Bridge motif participates in UL9 dimerization and likely contributes to allosteric regulation of UL9 function. This unique structural feature distinguishes UL9 from canonical SF2 helicases, particularly the DExD/H-box subfamily, suggesting that herpesvirus UL9-like helicases have evolved specific structural adaptations to support novel viral functions, potentially through novel molecular mechanisms distinct from those of cellular SF2 helicases.

### Structural details of UL9 C-terminal domain

The C-terminal domain (CTD) is structurally organized into three subdomains (Figs. 1 and 2a, b). **The first** (residues 536-747 and 800-833), comprising twelve α-helices and two antiparallel β-sheet pairs, constitutes the structural core of the CTD and mediates UL9 dimerization through extensive interactions with the opposing protomer. Based on its function, we designate it the **C-terminal dimerization domain (CDD)**. **The second subdomain** (residues 748-799) primarily comprises four α-helices. Structural homology analysis using the Dali server[31] revealed that this is the only region of the CTD with significant similarity to known structures. The top hits were all SAP (SAF-A/B, Acinus, PIAS) domains, including MANF[35] (PDB: 6HA7, Z=5.3, RMSD=2.7 Å), CDNF (PDB: 8QAJ, Z=4.6, RMSD=2.5 Å), PIAS1[36] (PDB: 1V66, Z=3.9, RMSD=2.1 Å), and so on (Figs. 2f and Supplementary Fig. 5). Notably, these matches exhibited <15% sequence identity despite structural conservation; for instance, the identity between UL9 residues 748-799 and the MANF SAP domain is only 12.8%. The SAP domain, named after three founding members SAF-A/B, Acinus and PIAS, represents a compact 35-residue module characterized by a conserved helix-turn-helix fold[37]. It is widely found in proteins involved in chromatin organization, transcriptional regulation, DNA repair, and RNA metabolism, and is well-characterized for its DNA/RNA-binding properties[37, 38] (Supplementary Fig. 5b). Given that this subdomain in UL9 functions as the determinant for dsDNA binding, we classify it as a **SAP domain[37]**. **The third region** comprises the final 17 residues (residues 834-851) located at the extreme C-terminus, which protrudes from the core CDD. Due to its flexible, tail-like conformation, we term this extension the C-terminal tail (**C-tail**). As detailed below, the C-tail facilitates UL9 dimerization, mediates ICP8 interactions, and modulates UL9 activity.

### Structural details of UL9 dimerization

Under physiological conditions, UL9 functions as a dimer[15, 29]. However, the specific dimerization mechanism has remained unclear due to limited structural information[29]. In this study, purified UL9 exists definitively as a dimer in solution, in both apo and DNA-bound states, as confirmed by cryo-EM (Figs. 1b, c and Supplementary Fig. 2). Moreover, DNA binding induces no observable conformational changes in the UL9 dimer structure (Supplementary Fig. 4c). Structural analysis reveals that UL9 homodimerization is mediated primarily by symmetric interactions between the CDD and C-tail of one protomer and the CDD and helicase domain of the opposing protomer, creating an overall interface area of approximately 3815 Å^2^. Notably, no direct interprotomer interactions were observed between the helicase domains, and the SAP domain is not involved in dimer formation.

Based on structural features, we classified the dimerization interface into three distinct types (Fig. 3). The **type I interface** constitutes the C2-symmetric dimer core formed by two adjacent CDDs (Fig. 3b). This interface is stabilized mainly by hydrophobic interactions, including the residues L552, A603, R606, H611, V616, L628, A630, P633, L635, A823, I826, L827, A830, L831, and L834 from two protomers. Additionally, E555 and D824 of one protomer form a salt bridge and a hydrogen bond with R607 of the other protomer, respectively, while E548 forms a hydrogen bond with the main-chain of A603 (Supplementary Fig. 6a). The **type II interface** is formed between the CDD of one protomer and the helicase domain of the opposing protomer (Fig. 3c). Residues P596, M597, and P835 of the CDD make van der Waals contacts with residues V410, F430, Y469, and C473 of the Bridge motif. Furthermore, residues T599, S602, E642, S832, and E833 of the CDD form salt bridges or hydrogen bonds with N373 of RecA2 and residues R406, S407, E408, P409, Q429 and Y469 of the Bridge motif, respectively (Supplementary Fig. 6b). The **type III interface** is established by the insertion of C-tail into the inter-domain cleft formed by the helicase domain and the CDD of the opposing protomer (Fig. 3d). C-tail residues P835, A838, W839, P840, M841, and M842 make van der Waals contacts with residues V350, V353, and P376 of RecA2, residues H468 and Y469 of Bridge, and residues A545, V550, V554, V562, P564, and I567 of the CDD. Additionally, C-tail residues T836, E837, A838, and Q843 form salt bridges or hydrogen bonds with residues R112, Y184, S185, and T187 of RecA1, N373 of RecA2, R472 of Bridge, and N820 of the CDD of the opposing protomer (Supplementary Fig. 6c). Structurally, the C-tail serves as a ‘molecular glue’ that simultaneously tightly bridges multiple subdomains (RecA1, RecA2, Bridge and CDD) of the opposing protomer and potentially inhibits inter-subdomain movement. This symmetric interaction likely stabilizes the UL9 dimer conformation and contributes to allosteric regulation of its enzymatic activity.

**Figure 3.**
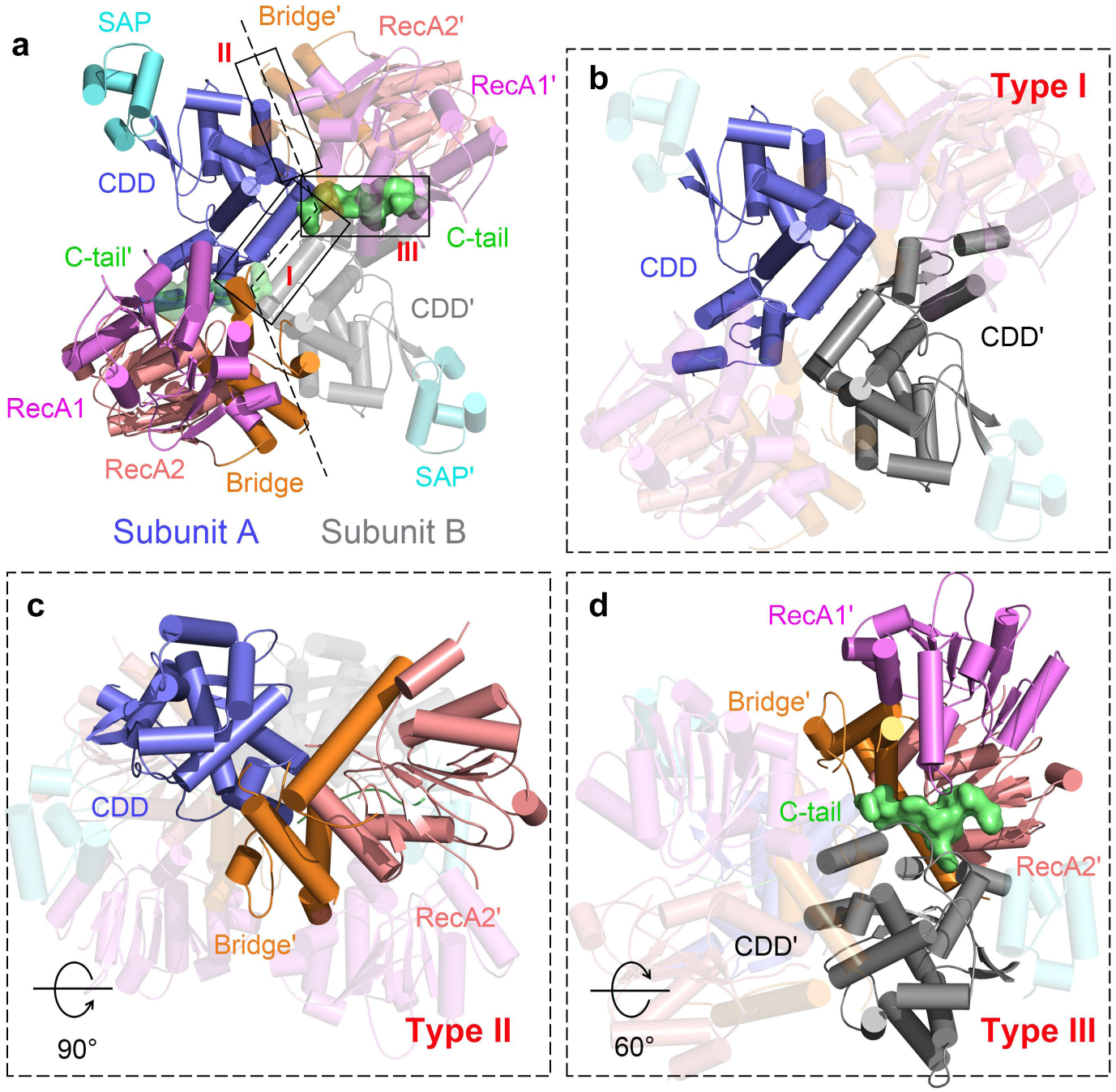
Structural details of UL9 dimerization. **a** Three types of dimerization interfaces classified in UL9. The two protomers are separated by a dashed line, with interfaces highlighted by solid rectangles and labeled. **b** Symmetrical Type I interface formed between CDDs. **c** Type II interface involving the CDD of one protomer and the RecA2 subdomain and Bridge motif of the opposing protomer. **d** Type III interface comprising the C-tail of one protomer and multiple subdomains (RecA1, RecA2, Bridge and CDD) of the opposing protomer. In panels **b**-**d**, subdomains involved in interactions are represented as cartoons with labels, whereas regions without interaction are shown as transparent cartoons.

Sequence alignment (Supplementary Figs. 7 and 8) indicates that interfaces I and II are broadly conserved across most analyzed alphaherpesviruses or maintain conserved biochemical properties. In contrast, interface III shows significantly higher conservation, with 11 identical and 7 similar residues. Among human-infecting herpesviruses, while interfaces I and II show high conservation within alpha or beta herpesviruses respectively, clear divergence exists between the two subfamilies. Interface III retains greater cross-subfamily conservation, preserving 7 identical and 5 similar residues. A critical structural difference is the absence of the C-tail in beta herpesviruses, despite its functional importance in alphaherpesviruses. These observations indicate that while the general dimerization strategy remains relatively conserved between alpha- and betaherpesviruses, significant divergence has occurred in mechanistic details, particularly regarding C-tail-mediated dimerization and functional regulation.

### Interaction details between UL9 and dsDNA

The HSV-1 genome contains three distinct origin sites for viral DNA replication[9, 39]. UL9 specifically recognizes and binds to these sequences to initiate DNA synthesis[13, 39, 40]. To elucidate the molecular mechanism of the UL9–origin interaction, we determined the cryo-EM structure of the UL9-DNA complex. Notably, the structure reveals that the UL9 dimer binds to dsDNA in a 2:2 stoichiometric ratio (Figs. 1b, c and Supplementary Figs. 9a, b). Compared with apo-UL9, DNA binding induces no significant global conformational changes, except for the ordering of the SAP domain (Supplementary Figs. 2, 4). The structure clearly reveals that dsDNA binding is mediated primarily by the SAP domain (Figs. 1b, c and Supplementary Figs. 9a, b). Intriguingly, although the helicase domain is predicted to possess nucleic acid-binding capability, no specific DNA interactions with this domain were detected in our structure.

Despite attempts to determine the sequence-specific recognition mechanism of UL9, the flexibility of the SAP–DNA interface limited the local resolution to approximately 4.5 Å, preventing the acquisition of interaction details at the atomic level. Rigid-body fitting of the AlphaFold-predicted model into the density map suggests that UL9 likely interacts with dsDNA through residues Q749, R756, K758, N759, K762, R783, Y786, M790, and K793 of the SAP subdomain and residues K746, K802 and R804 of the CDD (Fig. 4a and Supplementary Fig. 9a). Predicted interactions include ionic contacts with the phosphodiester backbone mediated by basic residues (Arg/Lys) (R756, K762, R783 and K793 of the SAP subdomain, and K746, K802 and R804 of the CDD), hydrogen bonds with nucleobases formed by Q749, N759, and K758, and van der Waals stacking interactions involving Y786 and M790 (Fig. 4b). To validate these interactions, we employed electrophoretic mobility shift assays (EMSA) for binding confirmation and microscale thermophoresis (MST) assays for quantitative affinity measurements. As DNA binding is mediated primarily by the SAP subdomain, all assays were performed using the C-terminal domain constructs of UL9 (UL9^CTD^ or UL9C) and its derived mutants. The dsDNA substrate used in EMSA and MST assays consisted of a 15-bp Box I sequence from OriS. The results of EMSA (Fig. 4c) demonstrated that deletion of the SAP domain (UL9C^ΔSAP^) or the triple mutant R756A/K758A/K762A (UL9C^mRKK^) completely abolished the UL9^CTD^-DNA interaction. Single-point mutants K746A, R756A, K758A, K762A, Y786A, and M790A exhibited significant binding attenuation, whereas the Q749A, N759A, R783A, K793A, K802A and R804A mutants showed moderate but significant reductions. MST assays (Supplementary Fig. 10) confirmed that wild-type UL9^CTD^ binds to dsDNA with a dissociation constant *(K*_d_) of 2.23 ± 0.42 μM. Consistent with EMSA, no binding was detected for either UL9C^ΔSAP^ or UL9C^mRKK^. Affinities for mutants R756A, Y786A, and M790A decreased by approximately 5-to 14-fold; mutants K746A, K758A, K762A, K802A, and R804A showed 2- to 4-fold reductions; and mutants Q749A, N759A, R783A, K793A exhibited 1.4- to 1.8-fold reductions. The binding ability of mutant K793A was almost unaffected due to its distal location within the DNA-binding interface. The results of the MST assays are consistent with those of EMSA, jointly identifying the residues involved in UL9-DNA interactions. Collectively, these data support a coordinated mechanism involving the SAP domain and three adjacent CDD residues (K746, K802 and R804). These elements form a stable, triangular SAP-CDD-dsDNA interaction geometry (Fig. 4b) that likely stabilizes the SAP domain, facilitating its visualization in the cryo-EM structure.

**Figure 4.**
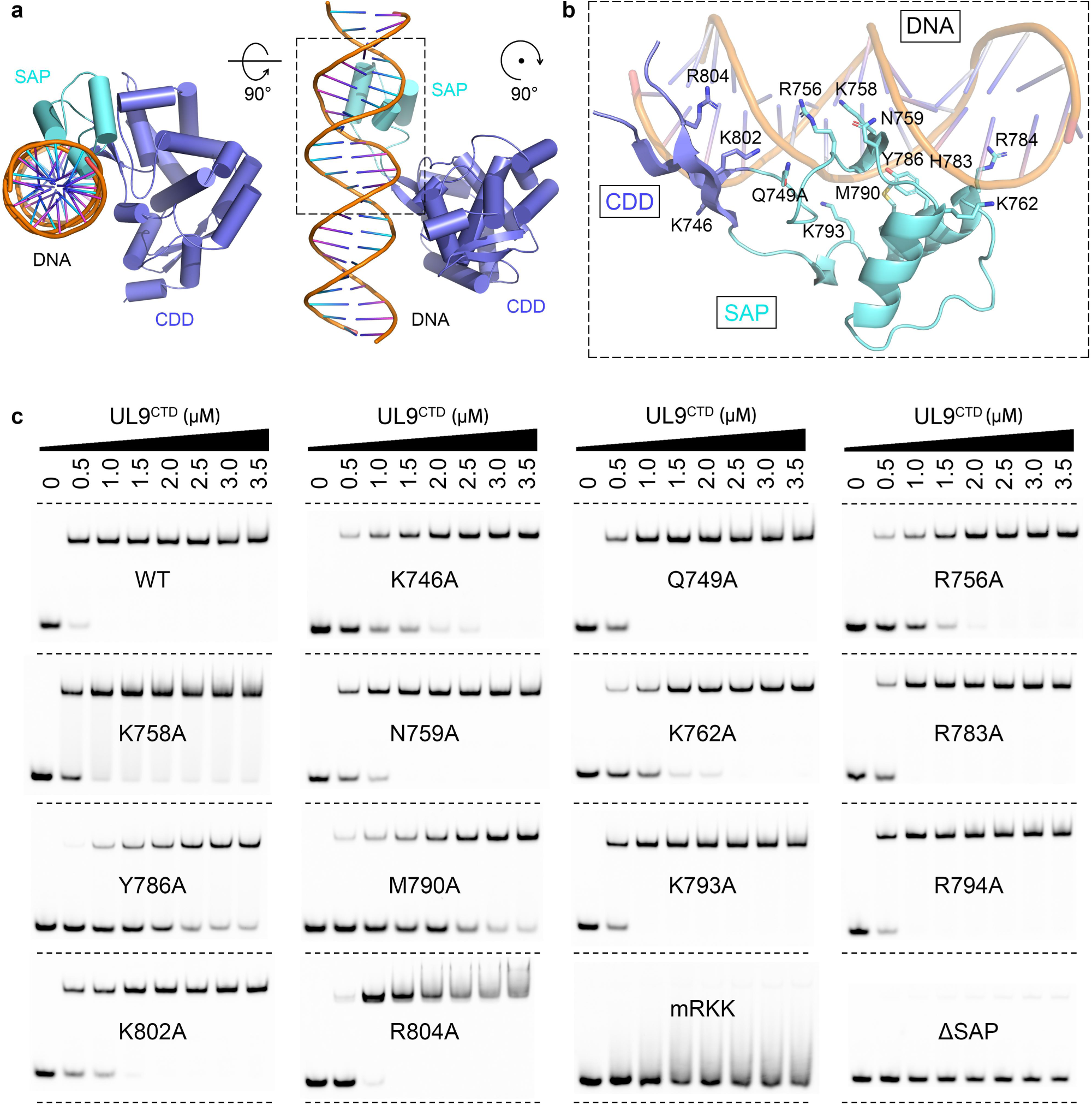
Details of the interaction between UL9 and DNA. **a** Two orthogonal views of the core structure of UL9-DNA interaction. **b** Structural details of UL9-DNA interaction. Panel **b** is an enlargement of the area enclosed by a dashed rectangle in panel **a** (dashed rectangle) after rotation. Protein and DNA structures are represented as cartoons, with side chains of residues involved in UL9-DNA interactions displayed as sticks. **c** EMSA analysis of wild-type UL9^CTD^ and its potential DNA-binding mutants. For EMSA assay, a series of seven increasing UL9^CTD^ concentrations (ranging from 0.5 to 3.5 μ M) were tested. All assays were independently repeated at least three times with consistent results.

Sequence alignment reveals remarkable conservation of DNA-binding residues across α-herpesviruses (Supplementary Fig. 7). Notably, five residues K758, N759, K762, Y786, and M790 are strictly conserved across all species examined, while four residues K746, R756, R783, and K793 consistently maintain positive charges (Arg or Lys). Residues Q749, K802, and R804 show limited variability, with functionally distinct substitutions observed in one or two of the ten analyzed species. Among human herpesvirus-encoded UL9-like proteins, distinct conservation patterns are revealed (Supplementary Fig. 8). Residues K762 and K802 are identical, and residues K758, R783, and K793 invariably carry positive charges. N759 shows high conservation in five out of six species, while other residues display clear divergence between α- and β-herpesvirus lineages.

### Structural basis underlying UL9-ICP8 interaction

UL9 interacts with the ssDNA-binding protein ICP8 to form a ternary replication initiation complex at the origin site[17, 41-43], and ICP8 stimulates the helicase activity of UL9[17, 19, 21, 42]. While previous studies have identified the C-terminus of UL9 as critical in ICP8 binding[17], the precise interaction site and mechanism remain unclear. To map the minimal ICP8-interaction region, we performed *in vitro* pull-down assays using recombinant UL9 truncation mutants guided by the structural information. First, a Strep-tag pull-down assay was employed to compare the binding capacity of UL9^FL^ and UL9^CTD^ for ICP8. The results showed that UL9^FL^ displayed markedly weak interaction with ICP8, whereas UL9^CTD^ exhibited strong binding (Fig. 5a). We next investigated the specific region of UL9^CTD^ participating in this interaction. GST pull-down assays showed that deletion of either the C-terminal 8 residues (UL9C^Δ8^) or 16 residues (UL9C^Δ16^) completely abrogated the UL9^CTD^-ICP8 interaction, whereas the isolated 8-residues (C-tail^8aa^) or 16-residues (C-tail^16aa^) segments showed no detectable ICP8-binding (Fig. 5b). Furthermore, deletion of the SAP domain did not impair the UL9C-ICP8 interaction, indicating that the SAP domain is dispensable for ICP8 binding. These results demonstrate that both the CDD and the C-tail are cooperatively required for effective ICP8-binding.

**Figure 5.**
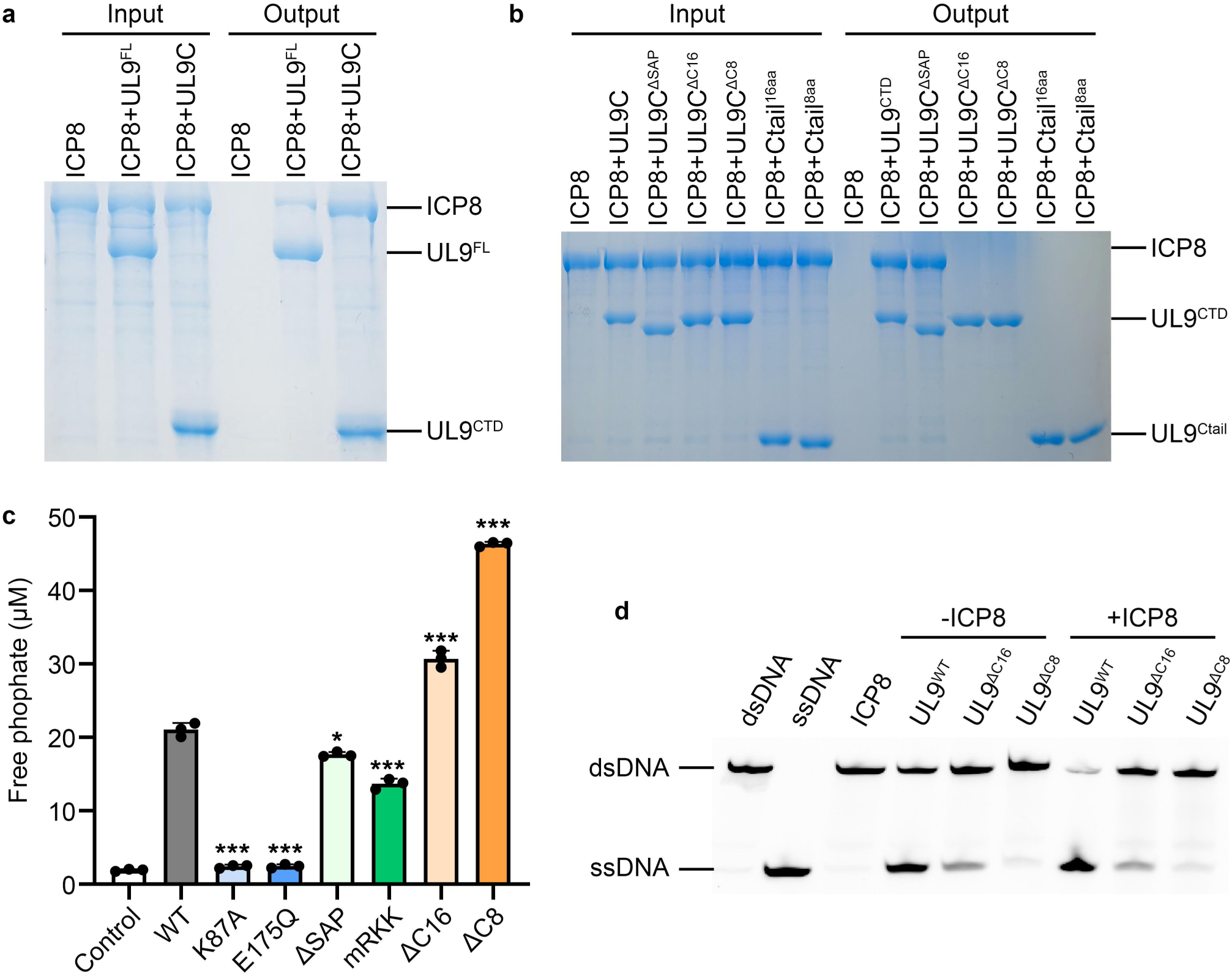
Interaction between UL9 and ICP8. **a** Strep-tag pull-down assay demonstrating interaction of ICP8 with either full-length UL9 or UL9^CTD^. **b** GST pull-down assay analyzing binding between ICP8 and either UL9^CTD^ or the C-tail. **c** Comparative ATPase activity of wild-type UL9 versus its functional mutants. ATPase activity assays were performed using the Malachite Green Phosphate Detection Kit. Data are presented as the mean with standard deviation for experiments performed in triplicate. Statistical significance was determined by a two-tailed Student’s *t*-test (*p < 0.05, **p < 0.01, *p < 0.001). **d** ICP8-mediated stimulation of UL9 helicase activity. Wild-type UL9 and two C-terminal truncation mutants (ΔC16 and ΔC8) were tested.

Integrating these findings with our structural data, we propose a mechanistic model for UL9-ICP8 interaction and its regulation. In the apo state, the interaction between UL9 and ICP8 is severely impaired because the C-tail is wrapped by the helicase domain of the opposing protomer, preventing cooperative engagement with the CDD for ICP8 interaction. In contrast, to enter the functional state, UL9 likely undergoes an ATP binding/hydrolysis-driven conformational rearrangement to release the C-tail, thereby permitting its cooperative engagement with the CDD for greatly enhanced ICP8 binding.

### Allosteric activation of UL9 activity by ICP8-binding

HSV-1 UL9 functions to unwind viral genome DNA at the replication origin through ATP hydrolysis-driven conformational changes, a process facilitated by ICP8 binding. Integrating the structural information, we systematically characterized the ATPase and helicase activities of UL9 and specific mutants under various conditions.

Purified recombinant wild-type UL9 exhibits both ATPase and helicase activities (Fig. 5c and Supplementary Fig. 11a). As expected, the ATP-binding-deficient mutant K87A and hydrolysis-deficient mutant E175Q completely lost both activities, confirming the essential role of ATP hydrolysis (Fig. 5c and Supplementary Fig. 11a). The DNA-binding-deficient mutants UL9^ΔSAP^ and UL9^mRKK^ retained basal ATPase activity but were completely defective in helicase activity due to loss of DNA-binding (Fig. 5c and Supplementary Fig. 11a). Furthermore, while DNA stimulates the ATPase activity of wild-type UL9, mutants UL9^ΔSAP^ and UL9^mRKK^ fail to respond to DNA stimulation (Supplementary Fig. 11b). These findings demonstrate that DNA-binding is required for helicase activity and stimulates ATP hydrolysis.

We next investigated the impact of the UL9 – ICP8 interaction on enzymatic activity. ATPase assays showed that ATP hydrolysis rate of UL9 increased in a dose-dependent manner with increasing ICP8 concentrations (Supplementary Fig. 11c). Compared to wild-type UL9, the C-terminal truncation mutants UL9^Δ8^ and UL9^Δ16^ exhibited constitutively enhanced ATPase activity, with UL9^Δ8^ exhibiting a 2.5-fold increase and UL9^Δ16^ a 1.5-fold increase (Fig. 5c), consistent with previous reports[40]. However, increasing ICP8 concentration did not further alter the ATPase activity of either mutant (Supplementary Fig. 11c). These results indicate that C-terminal truncation increases basal ATPase hydrolysis but abolishes ICP8-dependent regulation. In contrast to the enhancement of ATPase activity, C-terminal truncation progressively impaired helicase activity, with UL9^Δ8^ displaying more severe impairment than UL9^Δ16^ (Supplementary Fig. 11a). Moreover, while ICP8 markedly stimulated the helicase activity of wild-type UL9, neither truncation mutant responded to ICP8 stimulation (Fig. 5d).

Collectively, these results underscore the critical role of the C-tail in UL9 dimerization, ICP8 interactions and the allosteric regulation of both ATPase and helicase activities. These findings support the proposed role of the C-tail as a ‘molecular glue’ that bridges multiple subdomains of UL9, limiting inter-subdomain movement essential for coupling ATP hydrolysis to DNA unwinding. Because the C-tail interface on the helicase domain overlaps with a potential nucleic acid-binding site, the dynamic interaction between the C-tail and the helicase domain of the opposing protomer governs the modulation of both ATPase and helicase activities.

## Discussion

In summary, the cryo-EM structures of UL9 elucidate distinctive structural features and provide key functional insights. While the N-terminal helicase domain maintains the conserved RecA subdomains typical of SF2 helicases, UL9 possesses a unique, evolutionarily derived ‘Bridge motif’ that distinguishes it from other superfamily members. The C-terminal domain exhibits a clear structural division into three functionally specialized subdomains: (1) the CDD, which primarily mediates dimerization; (2) the SAP domain, responsible for sequence specific recognition of replication origin sites; and (3) the C-tail, which participates in both UL9 dimerization and ICP8 interactions, playing a crucial role in the allosteric regulation of enzymatic activity.

HSV-1 UL9 belongs to the herpesvirus-specific OBP family. As both an origin-binding protein and helicase, UL9 shares functional parallels to simian virus 40 (SV40) large tumor antigen (LTag)[44] and human papillomavirus (HPV) E1 protein[44]. However, unlike these SF3 helicases, which form hexamers[44, 45] via their helicase domains, UL9 functions as an obligate dimer mediated by its unique CDD rather than the helicase domain. Structural homology analysis confirms that the UL9 helicase domain most closely resembles the DExD/H-box SF2 helicase superfamily, with near-complete conservation of key residues involved in ATP binding and hydrolysis. Nevertheless, UL9 displays critical deviations. Unlike most SF2 helicases, which typically function as monomers[46, 47], UL9 is a dimer. Furthermore, the canonical Motif II sequence has undergone evolutionary divergence: distinct from DEAH in Prp2 and DECH in HCV NS3 (DEAH/D-box subfamily), DEAD in DDX48 and eIF4A (DEAD-box subfamily), DEVH in Ski2 (Ski2-like subfamily), and DQAD in Vasa (Vasa subfamily), HSV-1 UL9 contains 174-DEVM-177 (conserved in HSV-2 and VZV), while β-herpesviral homologs contain DEIM (HHV6A, HHV6B and HHV7) (Supplementary Fig. 8). Combined with the unique Bridge motif, these features clearly differentiate UL9 from well-characterized SF2 members. Given these distinct structural and mechanistic properties, alongside the conservation of dimerization and DNA recognition among herpesvirus OBPs, we propose classifying these proteins as a distinct ‘UL9-like’ subfamily within the SF2 helicase superfamily.

As an SF2 member, the UL9 helicase domain possesses potential nucleic acid-binding capability (Supplementary Fig. 9). While our structure reveals that primary dsDNA binding is mediated by the SAP domain, this configuration alone cannot fully explain the duplex DNA unwinding mechanism. We therefore propose that the helicase domain harbors a secondary DNA-binding site with specificity for ssDNA, as suggested by previous studies[48, 49]. This putative site likely cooperates with SAP domain to facilitate DNA unwinding via ATP hydrolysis-induced conformational changes.

During the preparation of this manuscript, Gustavsson et al.[50] reported cryo-EM structures of UL9–DNA complexes, including dimeric assemblies in the presence and absence of ATPγS, as well as low-resolution models of monomeric and tetrameric (dimer-of-dimer) forms. Their findings regarding UL9 dimerization, sequence-specific DNA binding, and the regulatory role of the C-terminal tail in modulating interactions with ICP8 and UL9 helicase activity align closely with our observations. However, whereas Gustavsson et al. proposed an asymmetric 2:1 (dimer:DNA) stoichiometry for the principal dimeric complex, our results robustly define a symmetric 2:2 complex (Supplementary Fig. 12). We attribute this difference arises primarily from variations in nucleic acid saturation during sample preparation. Our preparations employed a 1.5-fold molar excess of DNA over UL9 monomer to ensure stoichiometric saturation, whereas the prior study used a 1.2-fold excess. Given that the UL9 homodimer comprises two equivalent protomers, the simultaneous binding of two DNA duplexes is biochemically plausible *in vitro*. Re-inspection of the published density maps (EMD-52135, PDB 9HGI; EMD-52145, PDB 9HGJ) from Gustavsson et al.[50] at lower contour levels suggests the presence of residual, unmodeled density for a second DNA duplex (Supplementary Fig. 12). Furthermore, their reported tetrameric (dimer-of-dimer) assembly (EMD-52147) appears compatible with a 2:2 binding mode per constituent dimer (Supplementary Fig. 12). Mechanistically, a 2:2 stoichiometry enables a single dimer to bridge two distinct origin recognition boxes, providing a model whereby a single dimer can cooperatively coordinate multiple origin elements to drive DNA unwinding.

Beyond these structural comparisons, our work explicitly classifies the DNA-binding region as a SAP domain—a conserved eukaryotic fold characteristic of proteins involved in DNA and RNA metabolism. We validated the functional necessity of this domain through comprehensive EMSA and MST mutagenesis of critical residues. Crucially, our apo-state structure reveals that the SAP domain is intrinsically flexible, adopting an ordered conformation only upon DNA binding. This conformational plasticity likely promotes efficient sampling and capture of replication origins in solution, followed by stabilization through cooperative interactions with the CDD. Additionally, we investigated the potential function of the helicase domain as a secondary DNA-binding site that collaborates with the SAP domain to facilitate origin melting. In parallel, using a series of truncations, we probed possible allosteric modulation of UL9 activity by UL9 – ICP8 interactions. In summary, while both studies converge on the fundamental mechanisms of UL9 dimerization and DNA binding, the structural differences most likely represent distinct conformational snapshots captured under varying experimental conditions, thus offering complementary perspectives. Together, these complementary datasets provide a robust foundation for dissecting the molecular mechanisms underlying UL9-mediated replication initiation, which remains incompletely resolved and warrants further investigation.

Integrating the findings that ICP8 binds cooperatively to the UL9 C-tail and CDD to stimulate ATPase and helicase activities, we propose a four-stage **working model** (Fig. 6). In Stage I (Resting State), UL9 exists as a dimer with the C-tails inserted into the inter-domain clefts of the opposing protomers, maintaining the helicase domains in a low activity, auto-inhibited state. In Stage II (Activation), ICP8 binding extracts the C-tails from clefts, relieving auto-inhibition and activating the ATP hydrolysis. In Stage III (Unwinding), the activated helicase domains engage ssDNA and unwind the duplex DNA through ATP hydrolysis-driven conformational changes. In Stage IV (Termination), following unwinding, the ssDNA products are released and ICP8 dissociates from the complex.

**Figure 6.**
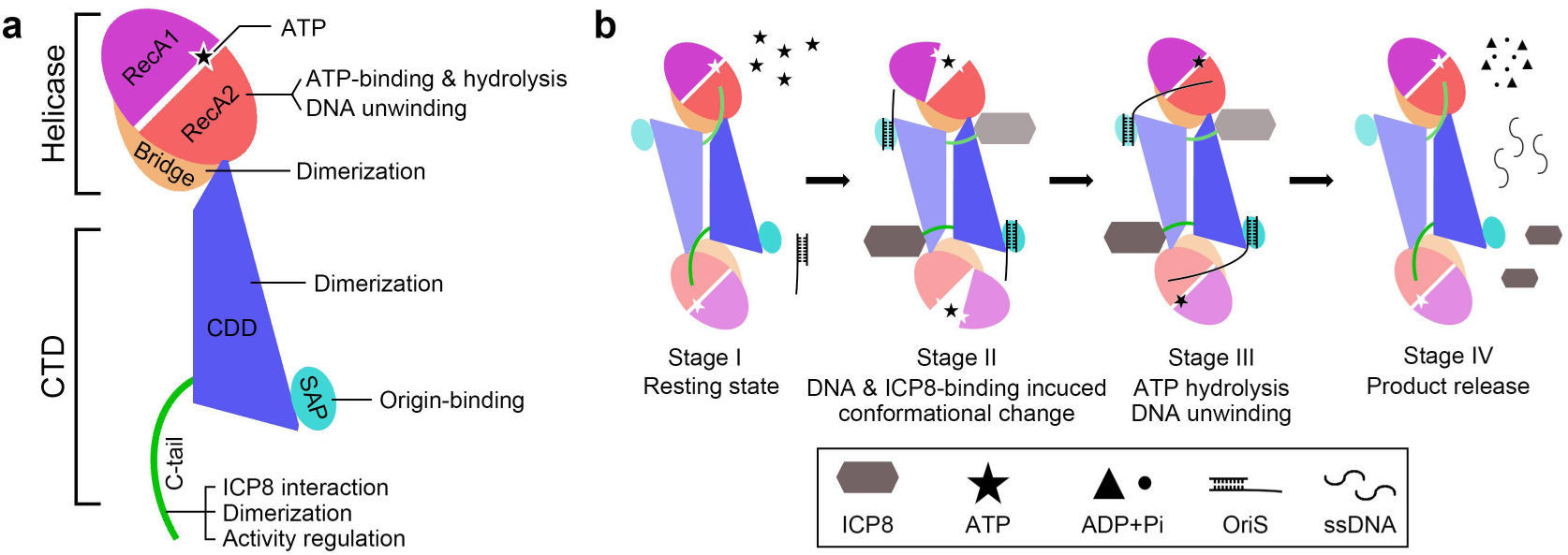
Working model of UL9 function. **a** A schematic diagram of HSV-1 UL9 showing domain organization and functional annotation. **b** Proposed catalytic cycle of UL9, divided into four stages. Stage 1, apo-UL9 dimer in resting conformation. Stage 2, conformational changes in UL9 dimeric complex induced by ICP8 and ATP binding, while dsDNA from the replication origin site binds to the SAP subdomain. Stage 3, coordinated ATP hydrolysis and ICP8 stimulation mediate dsDNA unwinding. Stage 4, release of reaction products (ADP and ssDNA), and ICP8 dissociation and recycling. Molecular component annotations are shown in the solid box below.

While this study defines the structural basis of UL9 dimerization, SAP-mediated origin recognition, and ICP8-dependent regulation, the precise molecular mechanisms underlying sequence-specific origin recognition and unwinding remain to be fully resolved. Future structural studies of UL9 complexes with origin DNA in distinct conformational states, as well as the UL9-ICP8-DNA ternary complex, are essential to fully elucidate these mechanisms. Furthermore, targeting the unique structural features of UL9 presents a promising avenue for the development of anti-herpesviral therapeutics.

## Materials and methods

### DNA substrates and preparation

All DNA substrates used in this study were synthesized by Tsingke Biotechnology and purified by HPLC. The oligonucleotides were initially dissolved in Annealing Buffer (10 mM Tris-HCl, pH 8.0, 50 mM NaCl, 1 mM EDTA) to a final concentration of 100 μM. For preparation of unlabeled dsDNA or fork DNA, complementary strands were mixed at a 1:1 molar ratio. For FAM-labeled constructs, the labeled strand and its unlabeled complementary strand were mixed at a 1:1.2 molar ratio. The mixtures were heated to 95 °C and denatured for 5 min in a thermal cycler (Bio-Rad), followed by gradually cooling to room temperature at least 3 h with the thermal cycler lid open. The annealed DNA products were stored at 4 °C and used within one week.

dsDNA substrate containing Box I and II from OriS used in cryo-EM single particle analysis: F_cryo-EM_: 5’-AAGCGTTCGCACTTCGTCCCAATATATATATATTATTAGGGCGAAGTGCGAGCACT-3’ R_cryo-EM_: 5’-AGTGCTCGCACTTCGCCCTAATAATATATATATATTGGGACGAAGTGCGAACGCTT-3’ dsDNA substrate containing Box I from OriS used in helicase and ATPase activity assay: F_Helicase_: 5’-CGCGAAGCGTTCGCACTTCGTCCTTTTTTTTTTTTTTT-3’ R_Helicase_: 5’-TTTTTTTTTTTTTTTGGACGAAGTGCGAACGCTTCGCG-FAM-3’ 15-bp dsDNA of Box I from OriS for MST assay and EMSA assay: F_MST_: 5’-GCGTTCGCACTTCGT-3’ R_MST_: 5’-ACGAAGTGCGAACGC-3’

### Protein expression and purification

The genes encoding full-length HSV-1 UL9 (UniProt: P10193) and ICP8 (UniProt: P04296) were codon-optimized and synthesized (GeneralBiol). UL9 was cloned into the pFastBac-1 (Invitrogen) vector with an N-terminal Strep-tag (^Strep^UL9) and expressed in Hi5 insect cells. After 48 h post-infection with P3 baculovirus generated in Sf9 cells, cells were harvested by centrifugation at 1500 × g for 10 min. The pellet was resuspended in Lysis Buffer (50 mM HEPES, 500 mM NaCl, 10% glycerol, pH 7.0) supplemented with cOmplete (Roche). The resuspended cells were sonicated for 15 min and centrifuged using a Beckman JA25.50 rotor at 20,000 × *g* for 40 min at 4 °C. The supernatant was retained and incubated with Strep-Tactin XT beads (IBA Lifesciences) for 1 h at 4 °C. The beads were then washed three times with Lysis Buffer to remove non-specifically bound proteins, and target protein was subsequently eluted using 2.5 mM D-biotin in Lysis Buffer. The target protein was pooled, supplemented with 1 mM DTT, concentrated (AmiconUltra, 50 kDa molecular mass cut-off) (Millipore), and further purified using Superdex 200 Increase 10/300 GL column (GE Healthcare) in Gel filtration buffer (20 mM HEPES, 300 mM NaCl, 1 mM DTT, pH 7.0).

The C-terminal domain of UL9 (residues 536-851, UL9^CTD^) was cloned into pET-22b vector (Novagen) with an N-terminal 8×His tag and overexpressed in *E. coli* BL21(DE3) (Novagen) cells. Protein expression was induced at 16 °C for 16-20 h with 0.3 mM isopropyl-1-thio-β-Dgalactopyranoside (IPTG) until OD600 ≈ 0.6. The harvested cells were resuspended in Lysis Buffer and lysed by three passes through a microfluidizer. After high-speed centrifugation at 20,000 × *g* for 40 min, the clarified supernatant was incubated with Ni-NTA agarose (GE Healthcare) for 1 h at 4 °C. Beads were washed three times with Lysis Buffer containing 60 mM imidazole, and target protein was eluted in Lysis Buffer containing 300 mM imidazole. The eluted protein was pooled, concentrated (10 kDa molecular mass cut-off), and further purified using Superdex 75 Increase 10/300 GL column (GE Healthcare) in Gel filtration buffer. All UL9^CTD^ mutants were purified using the same protocol as wild-type. UL9^CTD^ and its truncation mutants used for GST pull-down assays, along with two C-tail fragments, were cloned into the pGEX-6P-1 vector (GE Healthcare) to generate GST-tagged fusion proteins. Recombinant protein expression, cell resuspension, and lysis procedures followed the same protocol as for His-tagged UL9^CTD^. After centrifugation, the clarified supernatants were incubated with Glutathione-Sepharose 4B beads (GE Healthcare) at 4 °C for 1 h. The resin was then washed with Lysis buffer, and the GST-tagged target proteins were eluted using GST elution buffer (50 mM HEPES, 300 mM NaCl, 50 mM reduced glutathione, 1 mM DTT, pH 7.0). Finally, the eluted proteins were analyzed using SDS-PAGE, concentrated, and buffer-exchanged into Gel filtration buffer for subsequent analysis.

The full-length ICP8 was cloned into pFastBac-1 vector with an N-terminal 8×His tag and overexpressed in insect cells using the same protocol as UL9^FL^. Cell harvesting, resuspension, and lysis also followed the procedure outlined for UL9^FL^. ICP8 was purified via Ni-NTA affinity chromatography as UL9^CTD^, and then used for biochemical studies.

To prepare the UL9-dsDNA complex, purified UL9 and pre-treated dsDNA were mixed at a 1:1.5 molar ratio in 150 mM NaCl and incubated for 1 h on ice. The mixture was further purified using Superdex 200 Increase 10/300 GL column in buffer containing 20 mM HEPES, 150 mM NaCl, 1 mM DTT, pH 7.0 (Supplementary Fig. 1). Fractions containing the target complex were pooled and concentrated, and the fresh sample was used immediately for structural studies.

### Cryo-EM grid preparation and data collection

For cryo-EM grid preparation, freshly concentrated UL9 and UL9-dsDNA complex were first diluted to 0.6 mg ml^-1^ and 0.8 mg ml^-1^, respectively. The cryo-EM grids were prepared using an automatic plunge freezer Vitrobot Mark IV (ThermoFisher Scientific) under 100% humidity at 4 °C. Each aliquot of 3 μl diluted UL9 or UL9-dsDNA complex was loaded onto the freshly glow-discharged NiTi grid (R1.2/1.3, 300 mesh, Zhenjiang Lehua). After incubation for 10 s, excess sample was blotted with filter paper for 3 s with a blot force of 3, and then the grid was vitrified by flash plunging into liquid ethane. All datasets were automatically collected using SerialEM[51] through the beam-image shift data collection method[52] on a 300 kV Titan Krios microscope (ThermoFisher Scientific) equipped with an energy filter and a K2 Summit detector (Gatan). Images were recorded under the super-resolution mode at a nominal magnification of 165,000× and a defocus value ranged between -1.0 to -3.0 μm, resulting in a pixel size of 0.41 Å. Each move stack was exposed for 6.2 s with a total exposure dose of ∼60 electrons per Å^2^ over 32 frames.

### Data processing and reconstruction

All cryo-EM movie stacks were firstly binned and subjected to beam-induced motion correction and anisotropic magnification correction using MotionCor2[53]. Contrast transfer function (CTF) parameters were estimated with CTFFIND4.1[54] using non-dose-weighted images. Subsequent processing steps, including particle picking, 2D classification, 3D classification, and auto-refinement were performed in Relion[55] using dose-weighted images. First, particles were automatically picked using the Laplacian-of-Gaussian filter and processed with several rounds of reference-free 2D classification. Then, the best five images with clear structural features and diverse orientations were selected as templates for reference-based particle auto-picking. For apo-UL9, a total of 1,090,101 particles were picked from 2080 micrographs and subjected to reference-free 2D classification. 249,188 particles with clear structural features were kept for further 3D classification with C2 symmetry imposed, using the initial model generated by Relion. One among the five classes containing 158,949 particles was selected for final 3D auto-refinement, resulting in a 4.1 Å resolution density map (Supplementary Fig. 2). For UL9-DNA complex, 913,885 particles were picked from 2055 micrographs. After 2D classification, 147,241 particles were selected for further 3D classification with C2 symmetry, using the density map of apo-UL9 as the initial model. 110,702 particles from the best class among five were subjected to 3D auto-refinement and resulted in a 3.7 Å resolution density map (Supplementary Fig. 2). The local resolutions of final maps were calculated using ResMap[56]. Statistics for data collection and processing were summarized in Supplementary Table S1.

### Model building and refinement

For model building, the UL9-dsDNA complex was first built in Coot[57] with the help of the predicted model from Alphafold 2[30] based on its higher resolution and better quality density map. The cryo-EM density map of C-terminal domain (residues 536-844) in the UL9-dsDNA complex was good enough to trace almost all the residues; however, some density was missing at the N-terminal helicase domain (residues 19-535). The complete model was further improved by a few iterative rounds of real-space refinement using PHENIX[58] and manual modification in Coot until no further improvement could be achieved. The apo-UL9 model was generated by docking the refined model of UL9-dsDNA complex into the corresponding density map in UCSF ChimeraX[59], followed by iterative real-space refinement and manual modification. The final model of UL9-dsDNA complex includes residues 19-843 and two duplex DNA molecules, with residues 135-146, 267-283 and 431-456 missing. In contrast, residues 748-799 in the apo-UL9 model were not modeled due to the absence of corresponding density in the cryo-EM map. Statistics of the final models were summarized in Supplementary Table S1. Structural analysis was performed with UCSF ChimeraX and PyMOL (https://pymol.org/).

### ATPase activity assay

ATPase activity assays were performed using the Malachite Green Phosphate Detection Kit (Beyotime) according to the manufacturer’s instructions. The reactions were carried out by mixing 1 μM UL9 protein and 2.5 mM ATP of the final concentration in 20 μl ATPase Buffer (20 mM HEPES, pH 7.5, 150 mM NaCl, 5mM MgCl_2_, 1 mM DTT, 10% glycerol) in 96-well plates and incubated at 37 °C for 30 min. The reactions were terminated by adding 180 μl 200 mM EDTA, followed by addition of 70 μl malachite green reagent for colorimetric detection, and further incubated at room temperature for 30 min. The absorbance values of colorimetric products were measured at 630 nm using an Envison Multimode Microplate Reader (PerkinElmer). The phosphate standard curve was prepared according to the kit instructions. All assays were independently repeated at least three times.

### Helicase activity assay

FAM-labeled DNA substrates used for helicase activity assays were prepared by annealing a 3’-FAM-labeled DNA with an unlabeled complementary DNA at a molar ratio of 1:1.2. The reactions were carried out by mixing 50 nM dsDNA or fork DNA substrate with proteins of different concentrations in Helicase Buffer (20 mM HEPES, pH 7.5, 150 mM NaCl, 1 mM DTT, 10% glycerol, 5 mM MgCl_2_, 0.1% BSA, 2.5 mM ATP) and incubated at 37 °C for 30 min. The reactions were terminated by adding 1 mg/ml proteinase K for 20 min. The samples were analyzed using 15% Urea-PAGE and the images were taken using a ChemiDoc^TM^ MP Imaging System (Bio-Rad). All assays were independently repeated at least three times.

### Electrophoretic Mobility Shift Assay (EMSA)

DNA binding of UL9^CTD^ was analyzed using Electrophoretic Mobility Shift assay (EMSA). Briefly, 0.5-3.5 μM of purified UL9^CTD^ or its mutants were incubated with 40 nM FAM-labeled dsDNA for 1 h at 4 °C in 20 μl EMSA buffer (50 mM Tris, pH 8.0, 50 mM NaCl, 1 mM DTT). Then the samples were mixed with loading dye and analyzed by 7% non-denaturing PAGE gel in 1 × TG buffer (25 mM Tris, 192 mM glycine). The gels were imaged using a ChemiDoc^TM^ MP Imaging System (Bio-Rad) according to the manufacturer instructions. All assays were independently repeated at least three times.

### Microscale thermophoresis (MST) assay

The dsDNA binding affinity of UL9^CTD^ was performed using microscale thermophoresis (MST) assay on a Monolith NT.115 instrument (NanoTemper Technologies). Purified UL9^CTD^ or its mutants (Ligands) with serial dilution in MST buffer (50 mM Tris, pH 8.0, 150 mM NaCl, 1 mM DTT and 0.05% Tween-20) was mixed with 40 nM of FAM-labeled dsDNA (Target) as using in EMSA assay. The samples were loaded into premium coated capillaries (NanoTemper) after 30 min incubation in dark at room temperature. Measurements were performed at 25 °C with 20% LED power and Medium MST power with laser off/on times of 5 s and 10 s, respectively. Data were collected using MO.Control software (v3.0), and analyses using MO.Affinity Analysis 3 (v3.0) (NanoTemper) were performed by fitting the normalized fluorescence (F_norm_) versus protein concentration to a single-site binding model. All assays were independently repeated three times.

### Pull-down assay

For GST pull-down assays, 50 μg of purified GST-fused UL9^CTD^ mutants or C-tail peptides were mixed with an equal amount of ICP8. After a tenth of each mixture was taken out as input controls, 25 μl of prepared GST beads (GE Healthcare) and 1 ml Pull-down buffer (50 mM Tris, pH 8.0, 150 mM NaCl, 1 mM DTT and 0.05% Tween-20) were added to the remaining samples and incubated with gentle rotation for 1 h at 4 °C. The beads were then pelleted by centrifugation at 1,000 × *g* and washed three times with 1 ml Pull-down buffer, and subsequently analyzed using 12% SDS-PAGE gels stained with Coomassie blue. For Strep-tag pull-down assays, 50 μg of purified Strep-tagged UL9^CTD^ or full-length UL9 were mixed with an equal amount of ICP8. Subsequent steps were performed as described for the GST pull-down assay. All assays were independently repeated three times.

## Data availability

The cryo-EM density maps and corresponding atomic coordinates have been deposited in the Electron Microscopy Data Bank (EMDB) and Protein Data Bank (PDB), respectively: UL9 (PDB 9VJI, EMD-65114) and UL9-DNA complex (PDB 9VJH, EMD-65113). All data that support the findings of this study are available from the corresponding authors on reasonable request. Source data are provided with this paper.

## Acknowledgements

Cryo-EM data collection was carried out at the Center for Biological Imaging (CBI), Core Facilities for Protein Science at the Institute of Biophysics (IBP), Chinese Academy of Sciences (CAS). We thank Boling Zhu, Xujing Li and other staff members at the CBI for their support with data collection. We thank Mengsi Sun from the Biochemistry core of Shenzhen Bay Laboratory for technical support for MST analysis. The project was funded by the National Key R&D Program of China (2021YFA1301501, 2017YFA0504700), the National Natural Science Foundation of China (31930069), the Strategic Priority Research Program of the Chinese Academy of Sciences (XDB37040101), the Basic Research Program Based on Major Scientific Infrastructures, CAS (JZHKYPT-2021-05), and the Key Laboratory of Biomacromolecules, Chinese Academy of Sciences (ZGD- 2023-05). J.M. is supported by the Shenzhen Bay Laboratory Startup Fund (21330061) and the Beijing Nova Program (No. Z201100006820033).

## Author contributions

J.M. and X.Z. conceived the project and designed all the experiments. C.H., H.W. and J.S. performed protein expression and purification, and conducted biochemical and functional studies. C.H. prepared and screened the cryo-EM specimens. J.M. and C.H. collected and processed the cryo-EM data, and built the model. C.H., J.M. and X.Z. wrote the manuscript with input from all authors. All authors participated in experimental discussions, performed data analysis and interpretation, and approved the final version of the manuscript.

