## Supplemental Figures and Table 1 for "Structure and mechanism of the HSV-1 origin-binding protein UL9"

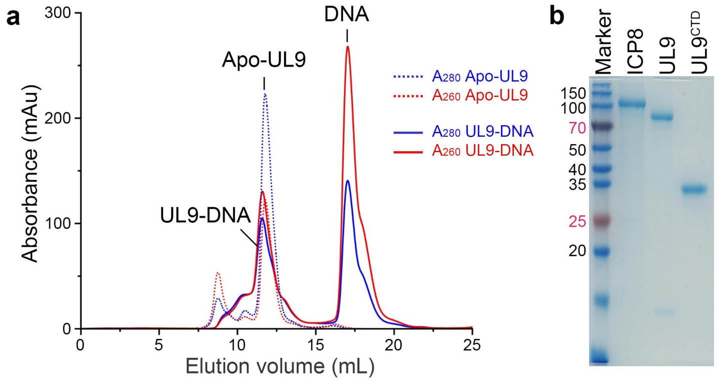


**Supplementary Figure 1. Preparation of recombinant HSV-1 full-length apo-UL9, UL9-DNA complex, UL9$^{CTD}$, and ICP8 proteins. a** Size-exclusion chromatography elution profiles of apo-UL9 and UL9-DNA complex using a Superdex 200 Increase 10/300 GL column. **b** Coomassie-stained SDS-PAGE analysis of purified UL9$^{FL}$, UL9$^{CTD}$, and ICP8 proteins. Molecular weights (in kilodaltons) of the marker are shown on the left lane, and bands are labeled above. The data shown are representative of at least three independent experiments.

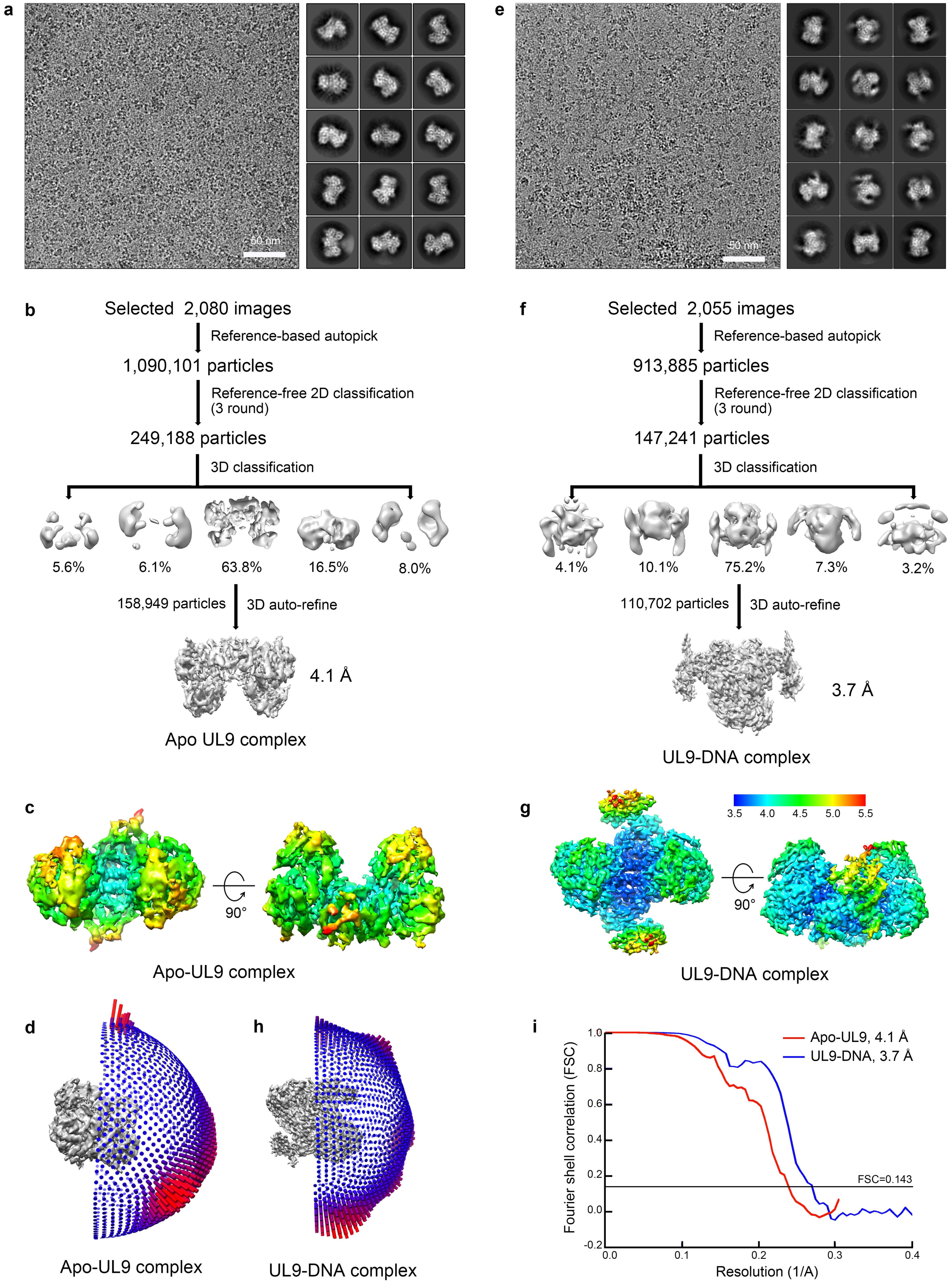


**Supplementary Figure 2. Single-particle cryo-EM analysis of HSV-1 apo-UL9 and UL9-DNA complex. a**, **e** Representative cryo-EM image and reference-free 2D-class averages of HSV-1 apo-UL9 complex (**a**) and UL9-DNA complex (**e**). **b**, **f** Data-processing workflow for apo-UL9 complex (**b**) and UL9-DNA complex (**f**). **c**, **g** Local resolution maps for apo-UL9 complex (**c**) and UL9-DNA complex (**g**) estimated by ResMap[56]. **d**, **h** Angular distribution of the particles of apo-UL9 complex (**d**) and UL9-DNA complex (**h**) in the final 3D auto-refinement. **i** The gold standard Fourier shell correlation (FSC) curves for the reconstruction of apo-UL9 complex (red line) and UL9-DNA complex (blue line) with resolutions at FSC = 0.143.

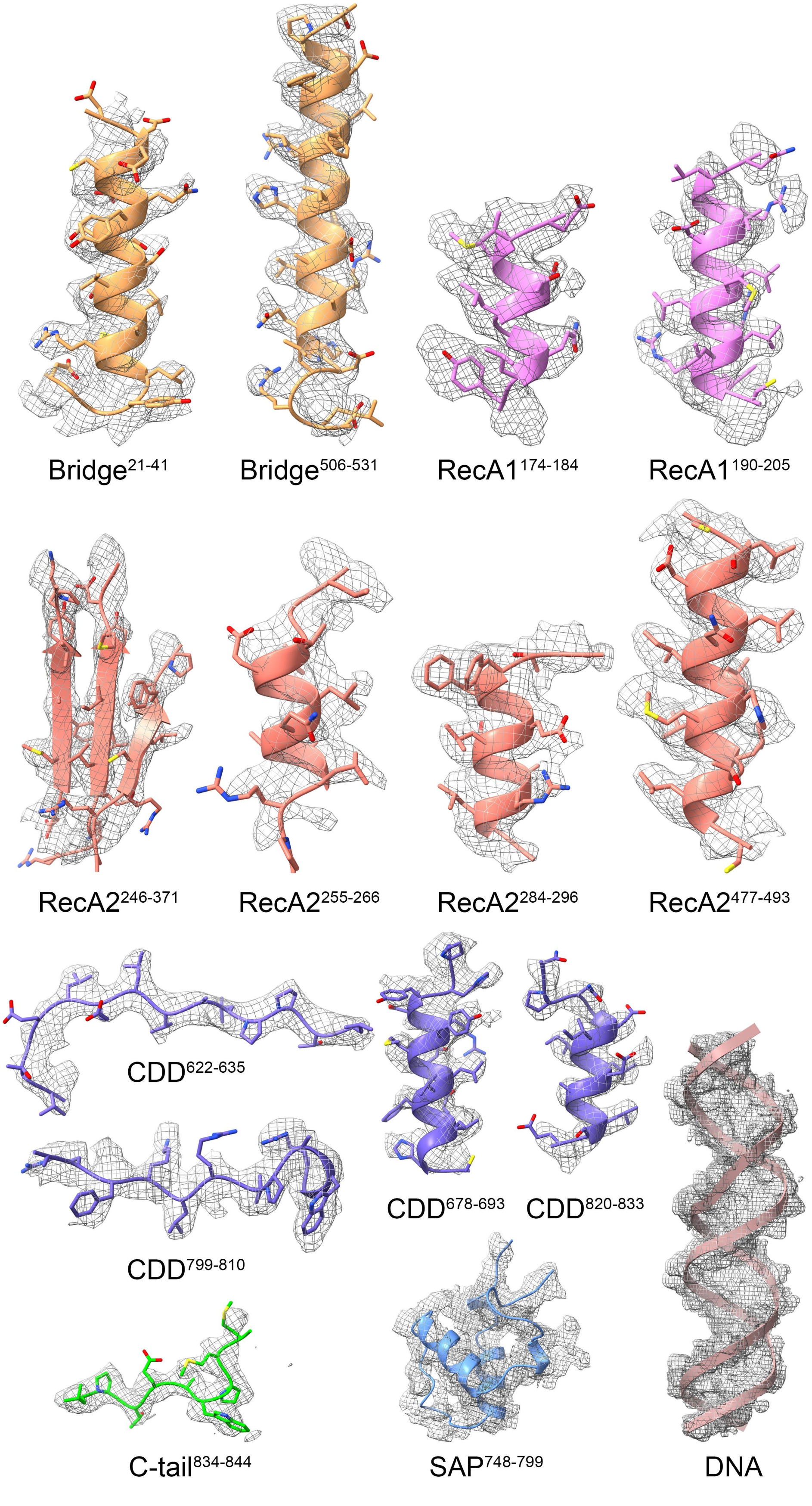


**Supplementary Figure 3. Representative density maps and corresponding atomic models of the UL9-DNA complex.** The densities are shown as gray meshes, while the corresponding models are shown as cartoons colored by domain as in Figure 1a, with side chains displayed as sticks.

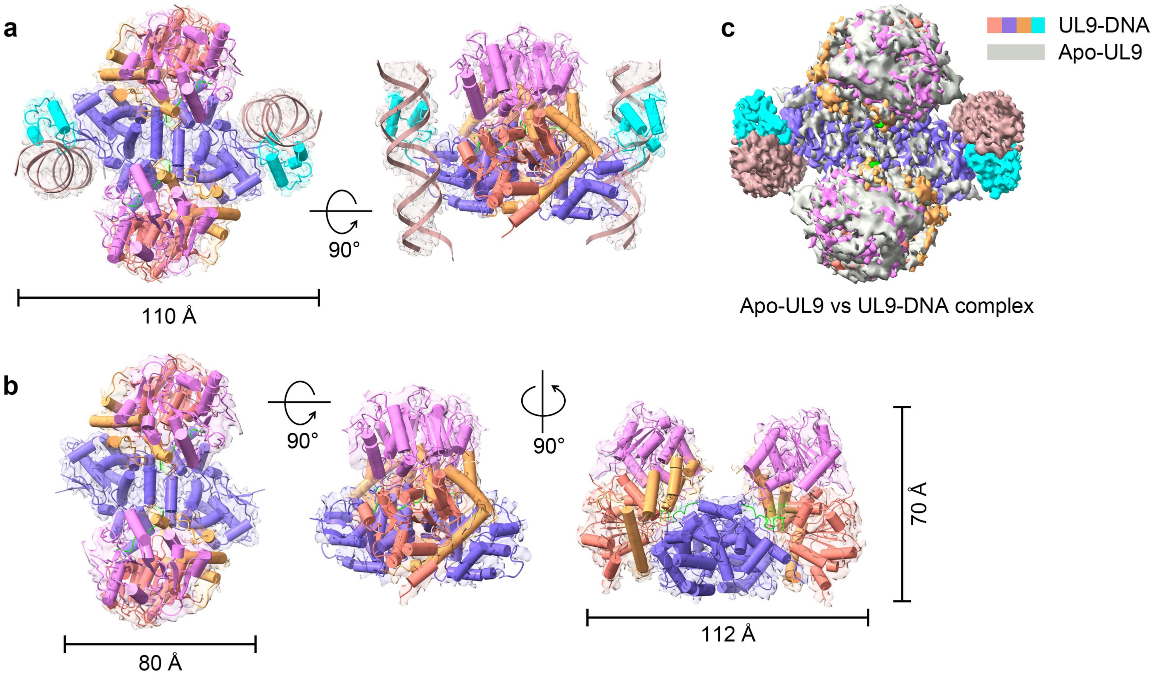


**Supplementary Figure 4. Structural comparison of apo-UL9 and DNA-bound UL9 complexes. a** Two orthogonal views of the UL9-DNA complex. **b** Three orthogonal views of the apo-UL9 complex. The density maps in panels **a** and **b** are displayed as transparent gray surfaces, with atomic models represented as cartoons colored by domain as in Figure 1a. The measured overall dimensions are labeled. **c** Superposition of cryo-EM density maps of apo-UL9 and UL9-DNA complexes. The UL9-DNA complex is colored by domain as in Figure 1a, while apo-UL9 complex is colored gray.

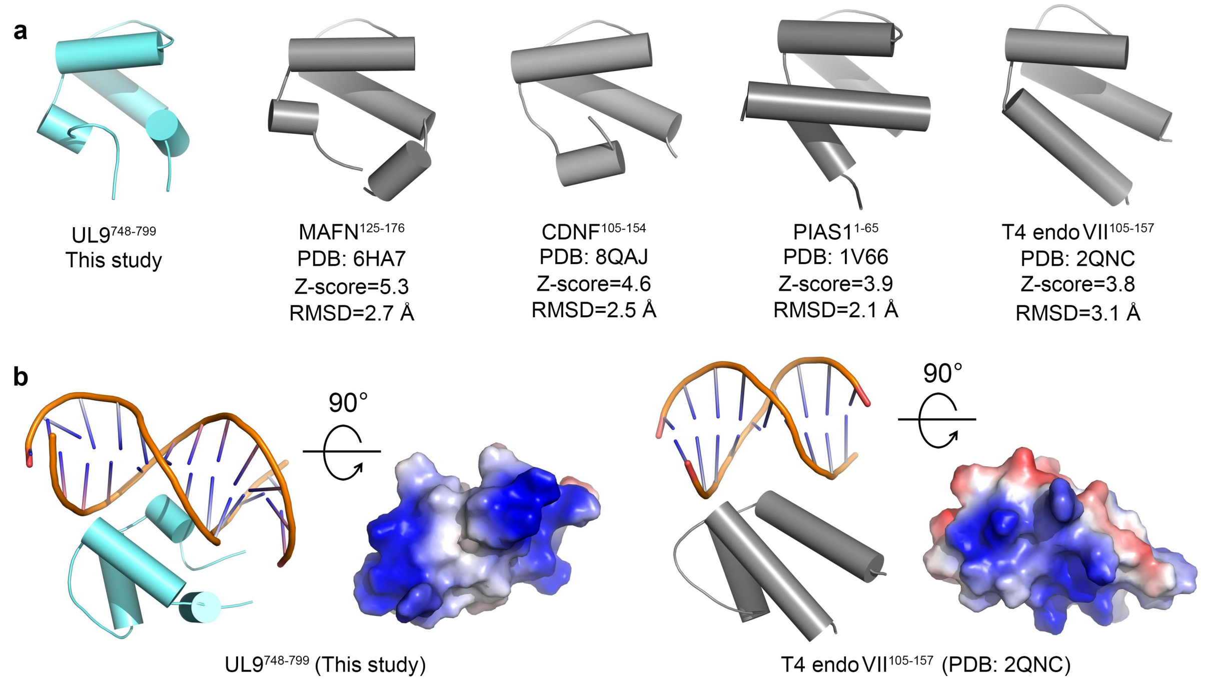


**Supplementary Figure 5. Homology analysis of the SAP domain of UL9. a** Comparison of SAP homologous structures searched through the Dali server. UL9 SAP is colored as in Fig. 1a, with homologous structures displayed in gray. Information about the homologs including PDB, residue range, Z-score, and RMSD is summarized below. **b** Comparison of DNA-binding features (DNA-binding model and surface electrostatic potential) between UL9 SAP and its homologous structure (T4 endo VII).

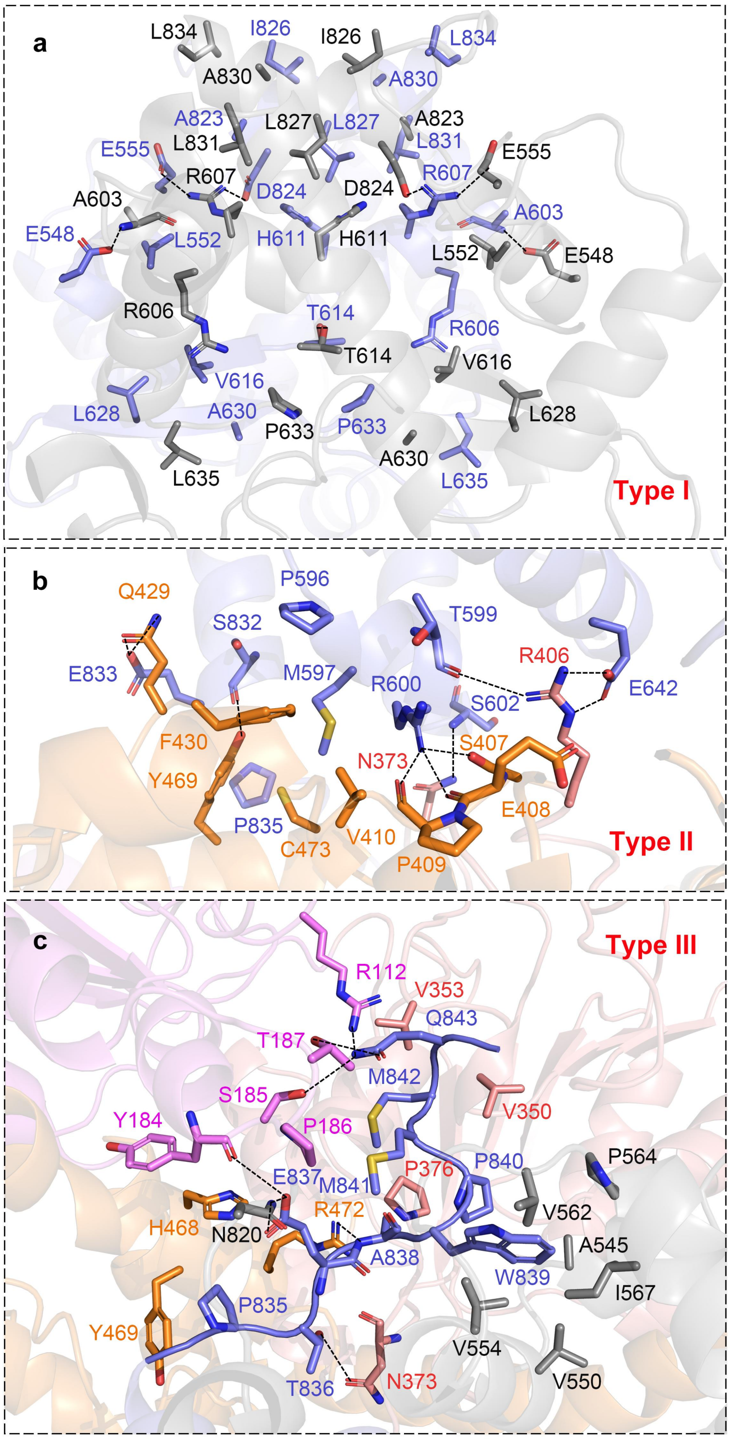


**Supplementary Figure 6. Structural details of UL9 dimerization interfaces.** The overall structure of the three distinct interfaces is shown in Figure 3, with the interaction details presented here. **a** Symmetrical Type I interface formed between the CDDs of two protomers. **b** Type II interface involving the CDD of one protomer and both Bridge motif and RecA2 subdomain of the other protomer. **c** Type III interface formed by the C-tail of one protomer interacting with multiple subdomains (RecA1, RecA2, Bridge and CDD) of the other protomer. The structures are represented as cartoons colored by domains as in Figure 1a, while side chains of residues participating in interactions are shown as sticks and appropriately labeled. Oxygen, nitrogen and sulfur atoms shown in red, blue and yellow, respectively, and salt bridges and hydrogen bonds are indicated as black dashed lines.

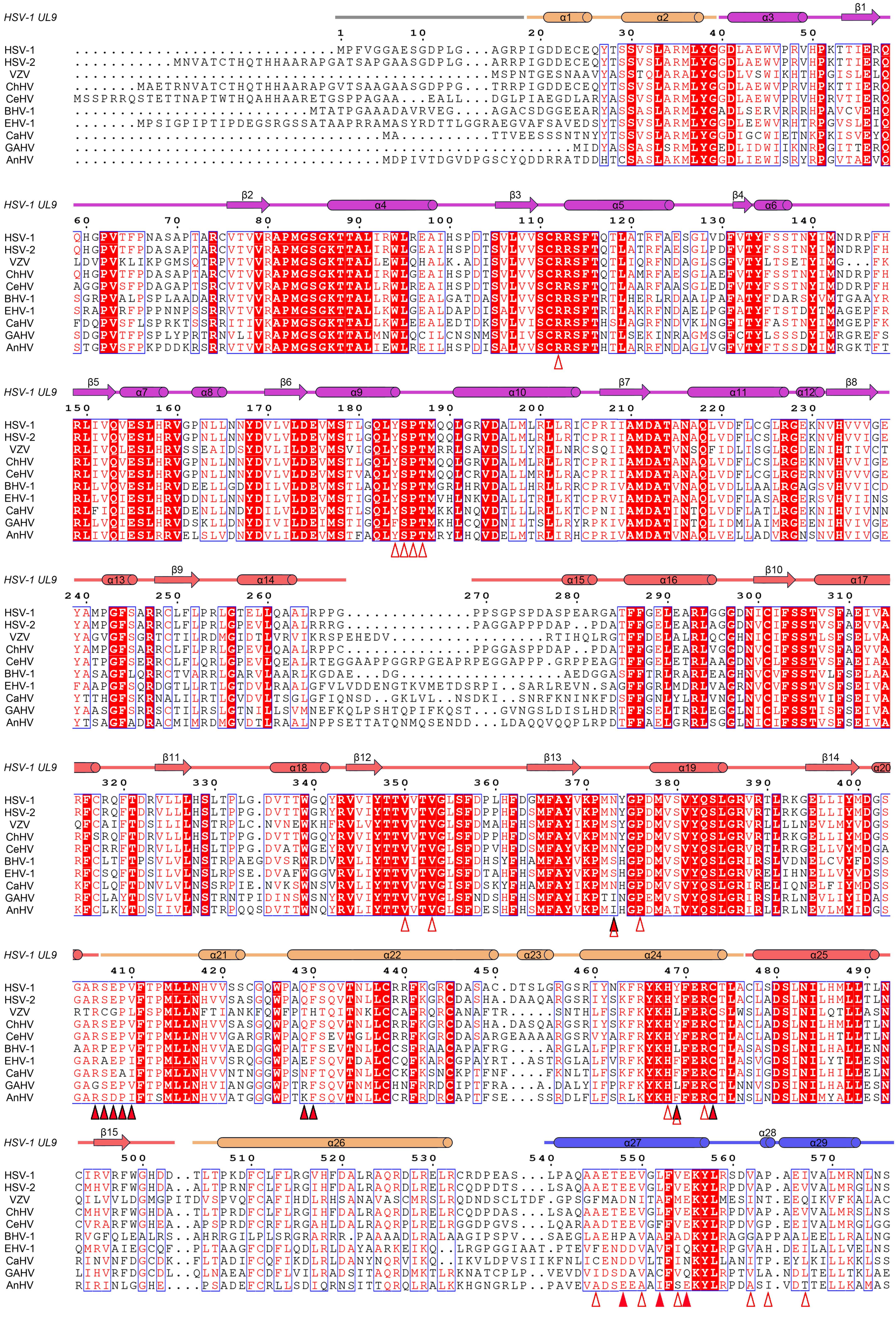
HSV-1 UL9
HSV-1
HSV-2
VZV
ChHV
CeHV
BHV-1
EHV-1
CaHV
GAHV
AnHV
α1 α2 α3 β1 β2 α4 β3 α5 β4 α6 β5 α7 α8 β6 α9 α10 β7 α11 α12 β8 α13 β9 α14 α15 α16 β10 α17 β11 α18 β12 β13 α19 β14 α20 α21 α22 α23 α24 α25 β15 α26 α27 α28 α29

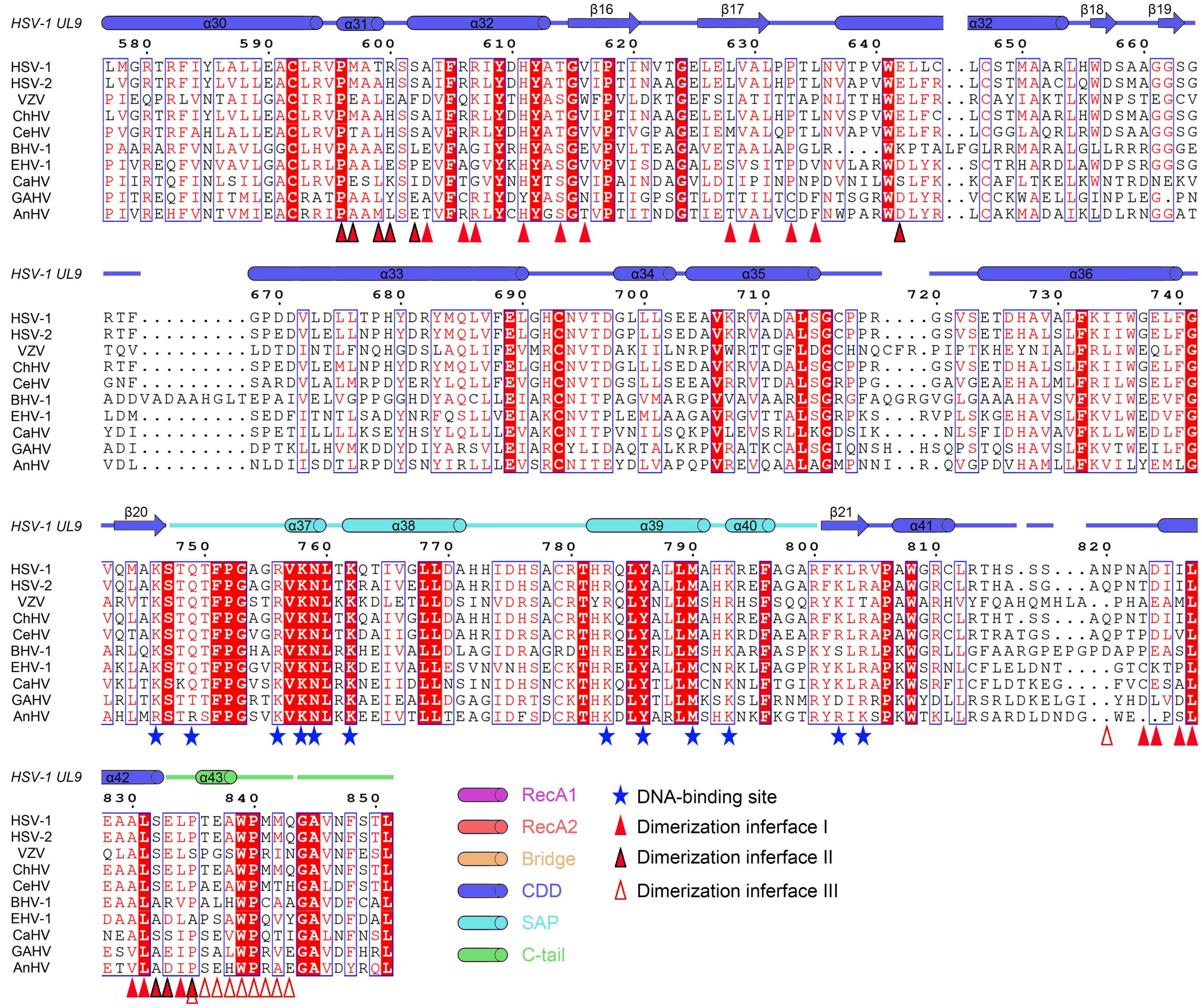

**Supplementary Figure 7. Multiple sequence alignment of representative α-herpesvirus UL9 homologs.** Uniprot accession numbers: HSV-1 (P10193), HSV-2 (P89432), VZV (Q9J3N7), ChHV (K9MG07), CeHV (X2FKV2), BHV-1 (P52377), EHV-1 (P84403), CaHV (A0A172ESJ6), GAHV-1 (Q9E6Q7), AnHV (A0A0U1YYV0). The structure-guided sequence alignment was generated using Clustal Omega[60] and rendered with ESPript[61]. Completely conserved residues are displayed as white text on red background, while similar residues are shown as red text on white background. Functionally important residues are annotated below the alignment. The solid red triangles, red triangles with black outline, and hollow red triangles represent dimerization interfaces I, II, and III, respectively, and blue asterisks denote DNA-binding residues. Secondary structure elements corresponding to the solved HSV-1 UL9 structure are depicted above the alignment, colored as in Figure 1a.

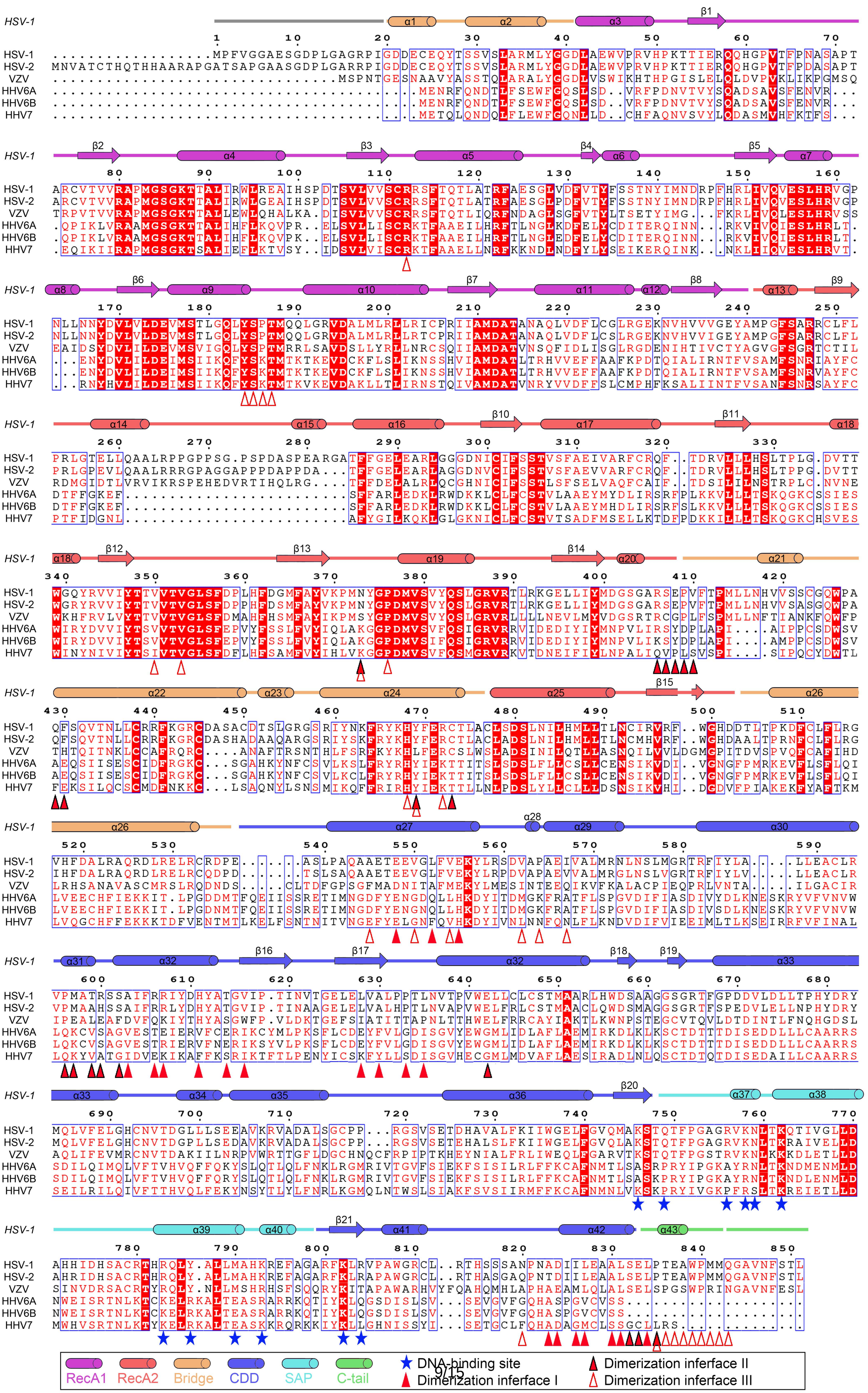
HSV-1
HSV-2
VZV
HHV6A
HHV6B
HHV7
DNA-binding site
Dimerization interface I
Dimerization interface II
Dimerization interface III
RecA1
RecA2
Bridge
CDD
SAP
C-tail

**Supplementary Figure 8. Multiple sequence alignment of the UL9-like proteins encoded by herpesviruses that infect human beings.** Uniprot accession numbers: HSV-1 (P10193), HSV-2 (P89432), VZV (Q9J3N7), HHV6A (P52378), HHV6B (P52452), and HHV7 (P52379). The generation of the sequence alignment and annotation of the results were performed as described for Supplementary Figure 7.

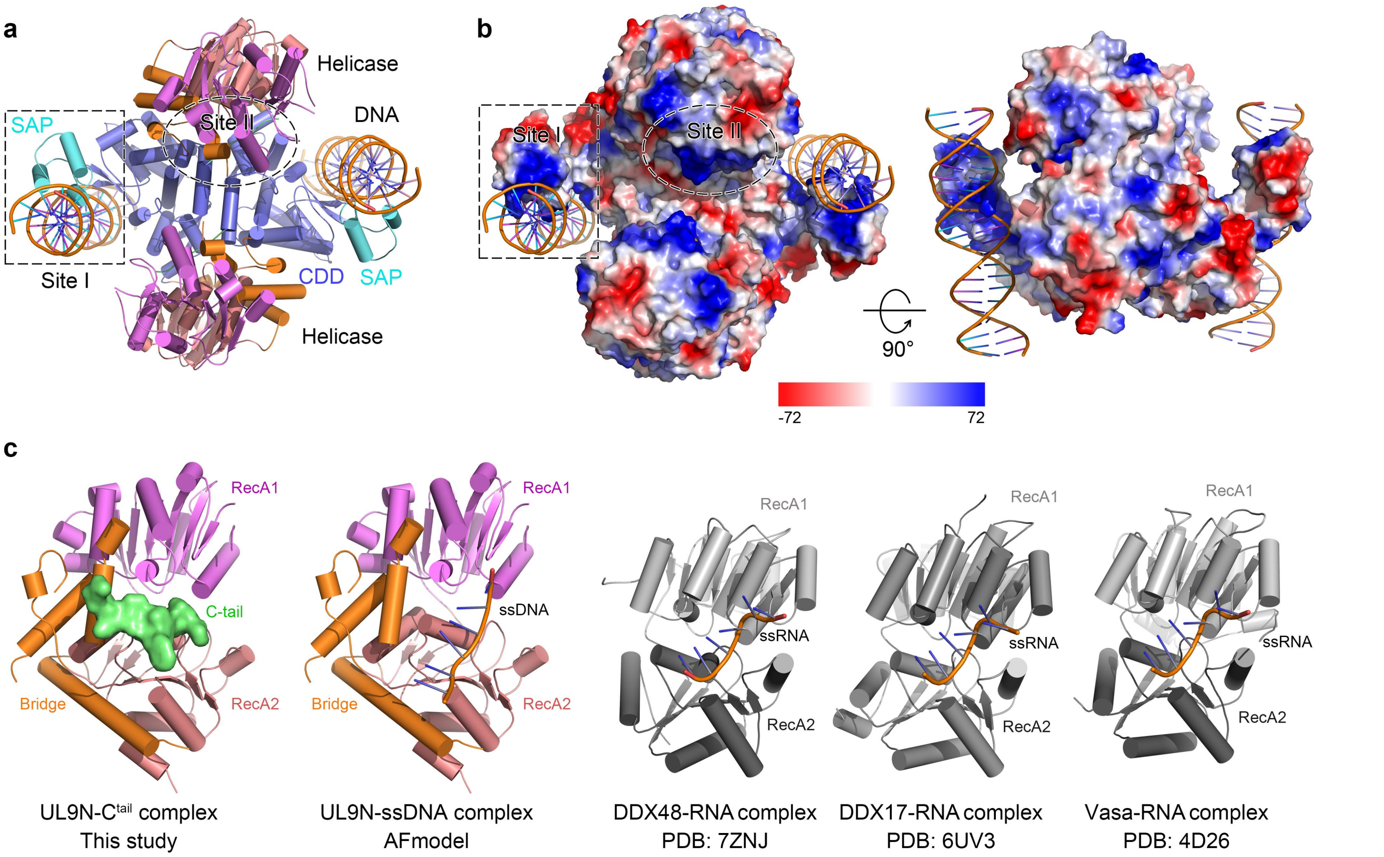


**Supplementary Figure 9. DNA binding properties of HSV-1 UL9. a** Structural characterization of UL9-DNA interactions shown in cartoon representation. DNA-binding Site I was identified in the cryo-EM structure of UL9-DNA complex. DNA-binding Site II represents a putative secondary DNA-binding region located on the helicase domain, but no direct helicase-DNA contacts were observed. **b** Two orthogonal views of surface electrostatic potential of the dimeric UL9 complex, with scale bar displayed below. **c** Structural comparison of UL9 helicase domain binding to the C-tail with a predicted UL9 helicase-ssDNA model from AlphaFold 3 and previous reported helicase-RNA complexes (DDX48-RNA, DDX17-RNA, Vasa-RNA). In panels (**a**) and (**b**), the solved Site I and predicted Site II are marked using dashed rectangles and ellipses, respectively.

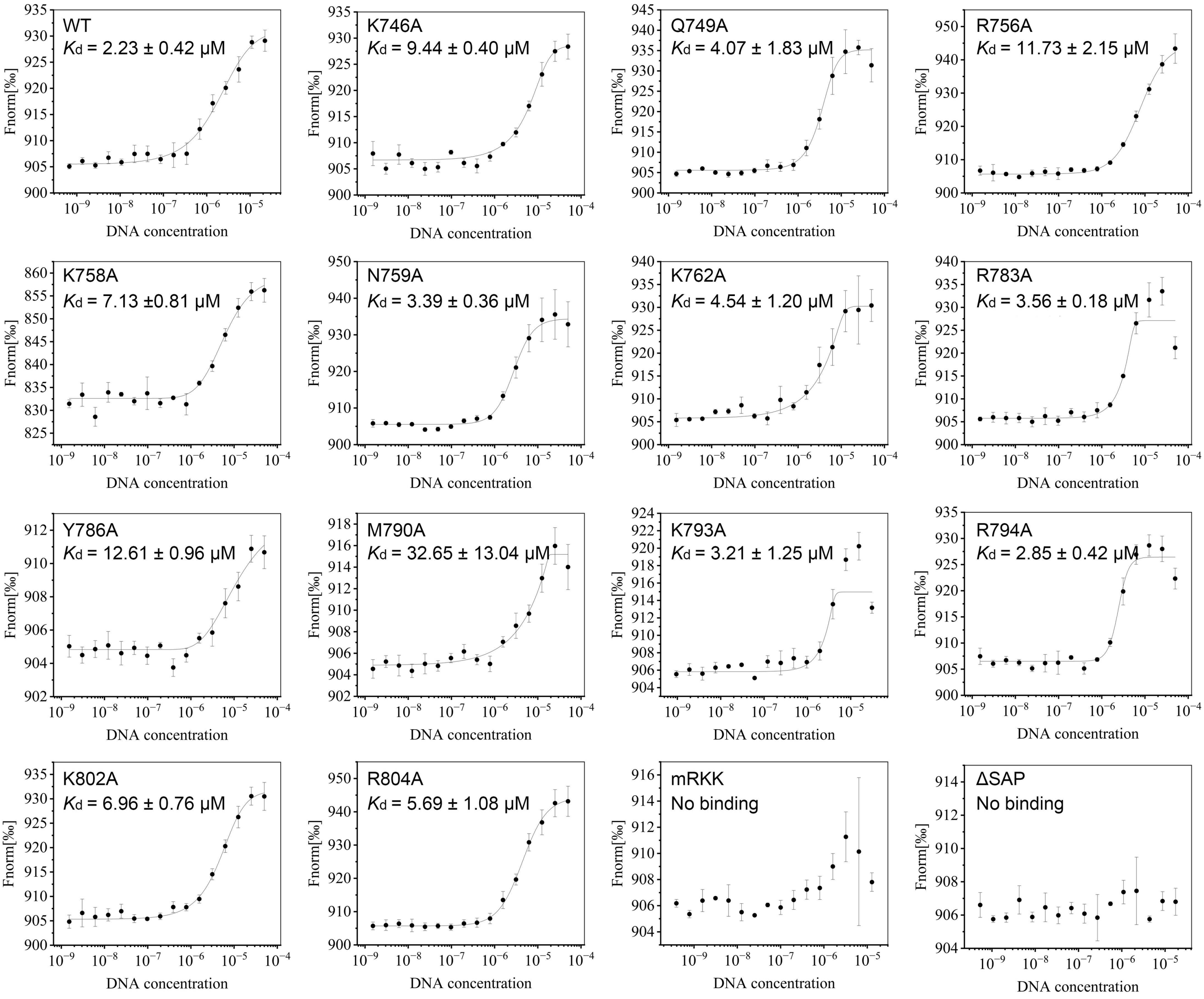


**Supplementary Figure 10. Microscale thermophoresis (MST) analysis of DNA binding by UL9$^{CTD}$ and its mutants.** The mutants and their corresponding dissociation constants ($K_d$) values are indicated in the upper-left corner of each MST assay profile. No binding indicates undetectable interaction between UL9$^{CTD}$ and DNA. All assays were repeated independently at least three times with consistent results.

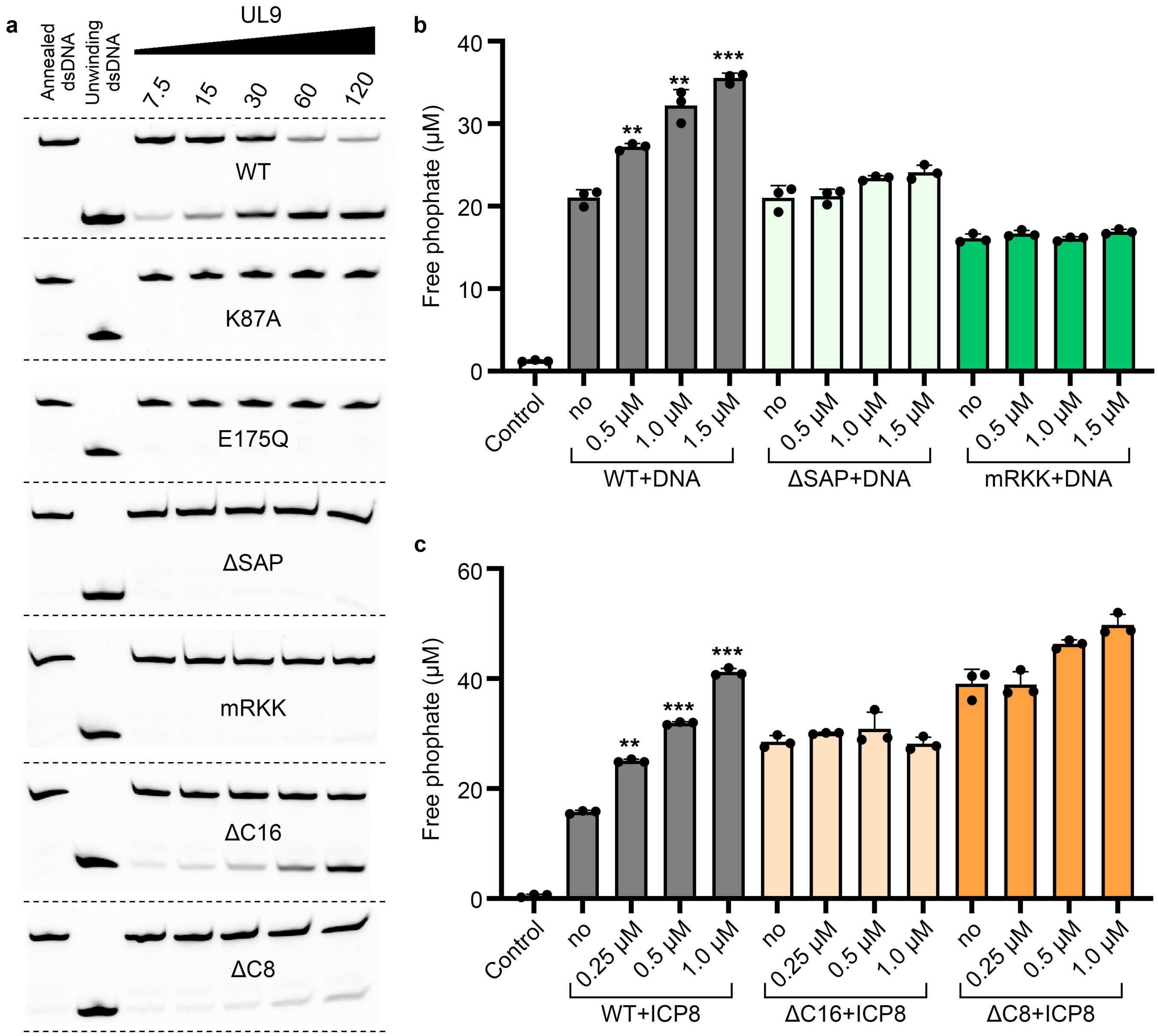


**Supplementary Figure 11. Comparative analysis of helicase and ATPase activities in wild-type UL9 and its mutants. a** Helicase activity of wild-type UL9 versus its functional mutants. The functional mutants include ATPase-deficient mutants (K87A and E175Q), DNA-binding-deficient mutants (SAP deletion ΔSAP and triple mutation mRKK), and ICP8-binding-defective C-terminal truncations (ΔC16 and ΔC8). Five different protein concentrations were used, ranging from 7.5 to 120 μM. **b** Comparative analysis of DNA-stimulated ATPase activity in wild-type UL9 versus DNA-binding-deficient mutants (ΔSAP and mRKK). **c** Comparative analysis of ICP8-mediated ATPase activity in wild-type UL9 versus C-terminal truncation mutants (ΔC16 and ΔC8). All assays were repeated independently at least three times with consistent results. Data represent mean ± s.d. of three independent replicates. P values were calculated by a two-tailed Student's *t*-test (*p < 0.05, **p < 0.01, *p < 0.001).

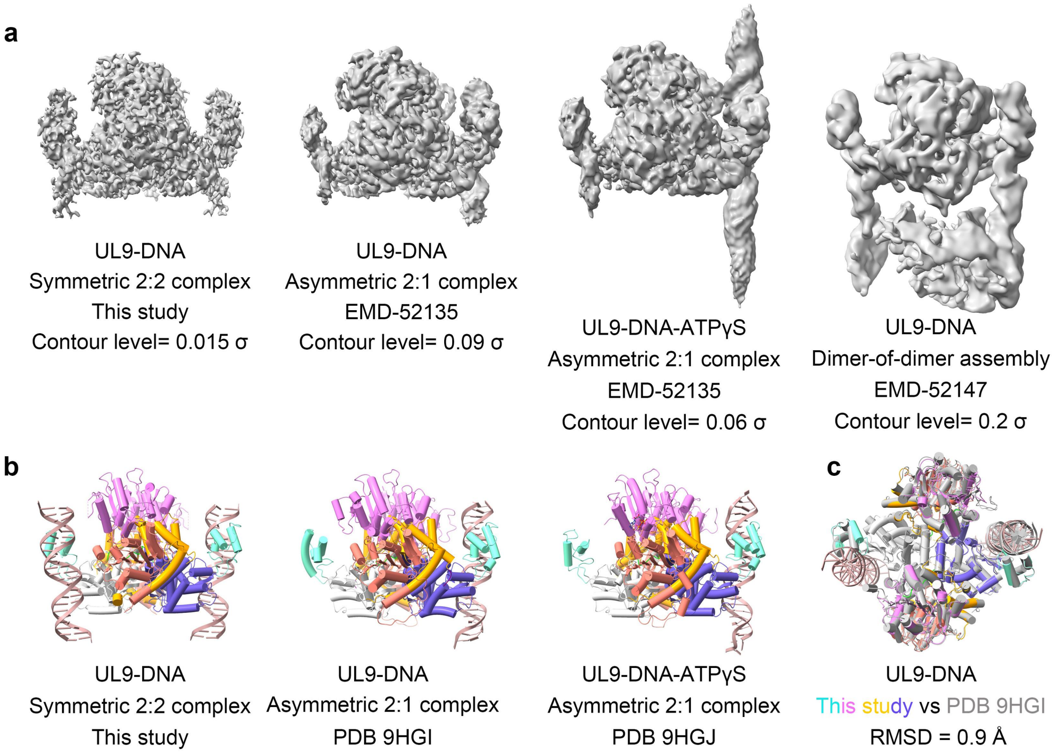


**Supplementary Figure 12. Structural comparison of UL9–DNA complexes. a** Comparison of cryo-EM density maps. The density map from this study is shown alongside those reported by Gustavsson et al.[50], including the dimeric assemblies in the presence and absence of ATPγS, and the tetrameric (dimer–dimer) assembly. EMD accession numbers and contour levels are indicated below each map. **b** Cartoon representations of UL9–DNA complexes. The structure determined here is displayed together with the dimeric structures reported by Gustavsson et al. in the presence and absence of ATPγS. Corresponding PDB accession codes are listed below each model. **c** Superposition of the UL9–DNA complex from this study onto the 2:1 UL9–DNA complex reported by Gustavsson et al. The Cα RMSD of 0.9 Å indicates near-identical overall architectures.

### Supplementary Table 1. Cryo-EM data collection and validation statistics.

| | Apo-UL9<br>EMDB-65114<br>PDB: 9VJI | UL9-DNA complex<br>EMD-65113<br>PDB: 9VJH |
|---|---|---|
| **Data collection and processing** | | |
| Microscope | FEI Titan Krios | FEI Titan Krios |
| Detector | Gatan K2 Summit | Gatan K2 Summit |
| Magnification (nominal/calibrated) | 165,000 | 165,000 |
| Voltage (kV) | 300 | 300 |
| Electron exposure ($e^-/Å^2$) | 60 | 60 |
| Defocus rang (μm) | 0.8~3.0 | 0.8~3.0 |
| Pixel size (Å) | 0.82 | 0.82 |
| Symmetry imposed | C2 | C2 |
| Initial particle images (no.) | 955,358 | 619,808 |
| Final particle images (no.) | 483,113 | 110,702 |
| Map resolution (Å) | 4.1 | 3.7 |
| FSC threshold | 0.143 | 0.143 |
| Map resolution range (Å) | 4.0~7.5 | 3.5~7.5 |
| **Model refinement and validation** | | |
| Initial model used | UL9-DNA complex | AlphaFold-predicted |
| Model resolution (Å) | 4.1 | 3.7 |
| Model resolution range (Å) | ∞~4.1 | ∞~3.7 |
| **Model composition** | | |
| Non-hydrogen atoms | 11226 | 13958 |
| Protein residues | 1432 | 1540 |
| Ligands | 0 | 4 |
| ***B* factors ($Å^2$)** | | |
| Protein | 47.34 | 49.41 |
| Ligand | | 131.11 |
| **R.m.s. deviations** | | |
| Bond lengths (Å) | 0.004 | 0.002 |
| Bond angles (°) | 0.934 | 0.523 |
| **Validation** | | |
| MolProbity score | 1.62 | 1.63 |
| Clash score | 8.45 | 8.19 |
| Rotamer outliers (%) | 0.00 | 0.00 |
| **Ramachandran plot** | | |
| Favored (%) | 97.09 | 96.91 |
| Allowed (%) | 2.91 | 3.09 |
| Disallowed (%) | 0.00 | 0.00 |
